# Synchronous spatio-temporal control of autophagy and organelle trafficking is necessary for appressorium-mediated plant infection by *Magnaporthe oryzae*

**DOI:** 10.1101/2025.10.06.680644

**Authors:** Alice B. Eseola, Xia Yan, Miriam Osès-Ruiz, Lauren S. Ryder, Martin J. Egan, Dan MacLean, Nicholas J. Talbot

## Abstract

The blast fungus *Magnaporthe oryzae* infects plants using a specialised infection structure called an appressorium that generates physical force to break the rice leaf cuticle. Appressorium development follows a cell cycle-controlled morphogenetic program, requiring autophagy-associated cell death of the fungal spore from which the infection cell develops. How proliferative growth of the fungus is regulated at the same time as programmed cell death, however, is unknown. In this study, we provide evidence that each cell of the conidium undergoes a separate developmental program, which is necessary for plant infection. Using quantitative live-cell imaging, we monitored trafficking of ten organelle types during appressorium morphogenesis in a wild-type *M. oryzae* strain and isogenic Δ*atg8* autophagic mutant. High-resolution microscopy using a photoactivatable green fluorescent protein revealed that organelle trafficking occurs from a single conidium cell into the appressorium, while the remaining two cells undergo autophagy. Organelle inheritance operates independently of cell cycle checkpoints but is always associated with spore germination. We furthermore defined the temporal sequence of organelle movement and *de novo* organelle biogenesis in the incipient appressorium using photoconvertible fluorescent localisation microscopy. Taken together, our study reveals how synchronous spatiotemporal control of autophagy and organelle trafficking is necessary for rice blast infection.

## Introduction

Many of the world’s most devastating plant diseases are caused by fungal pathogens that elaborate specialised infection structures called appressoria. Rusts, powdery mildews and anthracnose diseases for example, which collectively cause significant global crop losses (Savary et al., 2019), are all caused by appressorium-forming fungi. The importance of appressorium development is exemplified by the blast fungus *Magnaporthe oryzae* (also known as *Pyricularia oryzae*) (Zhang et al., 2016), where it is critical to its ability to infect numerous cereal hosts and cause the most serious disease of cultivated rice (Baudin et al., 2024; Ryder et al., 2022). The *M. oryzae* appressorium develops enormous turgor of up to 8.0 MPa which is applied as mechanical force at the leaf surface to break the cuticle and gain entry to plant tissue (Ryder et al., 2022). Plant infection is initiated when a three-celled spore of *M. oryzae*, called a conidium, attaches to the hydrophobic leaf surface using a powerful adhesive released from a periplasmic space at the spore apex (Osés-Ruiz et al., 2021). The conidium rapidly germinates to form a polarised germ tube, which hooks and swells at its tip, before differentiating into a dome-shaped melanised appressorium (Eseola et al., 2021).

Development of a functional appressorium requires three critical processes. First, the absence of external nutrients and exposure to a hard, hydrophobic surface triggers Pmk1 MAP kinase activation (Dean, 1997), which leads to phosphorylation of a large set of substrates, including a group of transcriptional regulators (Cruz-Mireles et al., 2024) that control appressorium formation (Osés-Ruiz et al., 2021; Sun et al., 2019). Secondly, cell cycle-dependent regulation is necessary, including an S-phase checkpoint that governs initiation of appressorium development, G2-M progression which controls appressorium maturation (Osés-Ruiz et al., 2017; Saunders et al., 2010a; Veneault-Fourrey et al., 2006a), and a pressure-dependent S-phase checkpoint required for septin-mediated re-polarisation of the appressorium (Osés-Ruiz et al., 2017). Cell cycle-dependent infection is mirrored in other appressorium-forming pathogens including barley powdery mildew *Blumeria graminis* (Hansjakob et al., 2012) and corn smut *Ustillago maydis* (Castanheira et al., 2014; Mielnichuk et al., 2009). Finally, the three-celled-conidium must recycle its contents through autophagy-dependent programmed cell death (Kershaw and Talbot, 2009; Veneault-Fourrey et al., 2006a), which may involve ferroptosis (Shen et al., 2020). Mutants impaired in autophagy form non-functional appressoria and are unable to cause rice blast disease (Kershaw and Talbot, 2009; Veneault-Fourrey et al., 2006a). Only if all these requirements are met will rice blast infection proceed.

These observations lead to an important series of questions regarding appressorium development. How does the blast fungus form a proliferative, actively growing cell– the appressorium –at the same time as undergoing regulated cell death? If a three-celled conidium is undergoing cell death, for example, how does organelle inheritance function to enable a functional pressurised appressorium to develop? And how are organelles and cytoplasm partitioned to facilitate fungal growth or broken down via autophagy to fuel appressorium turgor generation? When cells divide, for example, organelles must be partitioned between daughter cells, because many organelles cannot be readily made *de novo*, and instead must be distributed into newly formed cells where they act as templates for subsequent organelle biogenesis (Jorgensen et al., 2007; Warren and Wickner, 1996). Polarised cell growth offers a unique challenge to cellular organisation, because organelle inheritance requires specific trafficking processes to direct organelles into new formed cellular structures (Bowman, 2025). Our current knowledge of organelle inheritance, however, is based largely on studies in the budding yeast, *Saccharomyces cerevisiae*, where a preferred sequence of organelle movement occurs during bud emergence with endoplasmic reticulum (ER) and peroxisomes inherited ahead of vacuoles, mitochondria and, finally, nuclei (Li et al., 2021). In yeast, organelle movements are regulated independently of direct cell cycle control but are dependent upon bud emergence which precedes cell division. In other polarised cells, such as axon-containing neurons, relatively little is known regarding organelle distribution even though impairing these processes leads to neurological disorders (Suarez-Rivero et al., 2017), while in fungal hyphae organelles can move very long distances between cells but how organelle distribution is regulated is not well understood (Bowman, 2025).

In this study, we set out to investigate how trafficking of membrane-bound organelles occurs during appressorium development by the blast fungus and how this is orchestrated with the onset of autophagy. In this way, we sought to address how the fungus is able to develop a fully functional, actively growing differentiated infection cell at the same time as regulated cell death. To do this, we employed the use of high-resolution live-cell imaging to monitor the spatial dynamics of fluorescently tagged proteins over time (Rogers et al., 2021). Application of such 4D fluorescence live-cell imaging technique has been used in *S. cerevisiae* to document the sequential and spatial inheritance of organelles from mother to bud cells during cell division (Li et al., 2021). We also utilised conditional photoactivatable green fluorescent protein (paGFP) (Patterson and Lippincott-Schwartz, 2002) and the photoconvertible fluorescence mEos3.2 protein (Wiedenmann et al., 2004; Zhang et al., 2012) to functionally dissect the cellular control mechanisms governing appressorium morphogenesis.

We report that individual cells of the three-celled conidium do not contribute equally to providing cytoplasm and organelles to the developing appressorium. Instead, we show that two of the cells undergo *in situ* autophagy of their organellar contents, while the germinating cell is the single reservoir to provide cellular contents to the appressorium. Spatio-temporal control of organelle trafficking and *de novo* organelle biogenesis occur independently of cell cycle progression and autophagy, while proceeding in perfect synchrony. This reveals how location-and time-specific regulation of fungal autophagy and cellular trafficking are necessary pre-requisites for plant infection by the blast fungus.

## Results

### Subcellular localisation of specific organelles during appressorium development by *M. oryzae*

To investigate the spatio-temporal dynamics of organelle trafficking during appressorium development, we first identified markers to enable visualisation of nuclei, nucleoli, mitochondria, ribosomes, the endoplasmic reticulum, peroxisomes, plasma membrane, early Golgi bodies, the *trans*-Golgi network and vacuoles in *M. oryzae* using live cell imaging. To visualise their subcellular localisation, we generated C-terminal translational GFP fusions of each predicted organelle resident protein, expressed from its native promoter. We introduced each gene fusion into a wild-type strain of *M. oryzae* Guy11 and analysed transformants by live-cell imaging. All strains used in this study contain single GFP insertions and details of all strains generated in this study are summarised in Supplementary Table S1. The sub-cellular localisation of each organelle in conidia of *M. oryzae* is shown in Fig. 1. The nuclear type1 histone, H1-GFP, (Veneault-Fourrey et al., 2006a) localised to the nucleus of each cell of the conidium (Fig. 1A), confirmed by co-staining with DAPI (Supplementary Fig. S1A), and consistent with previous studies (Freitag et al., 2004; Kilaru et al., 2017; Veneault-Fourrey et al., 2006a). The nucleolar marker, Nop1-GFP, localised to a single compartment within the H1-RFP-labelled nucleus (Fig. 1B), (Saunders et al., 2010b) (Supplementary Fig. 1B), consistent with localisation reported for the nucleolar *Aspergillus nidulans* fibrillarin An-Fib (Ukil et al., 2009). The 60S ribosome marker, Rpl25-GFP, predominantly localised in the cytoplasm (Fig.1C), consistent with 60S ribosomal subunit localisation in *S. cerevisiae* (Gadal et al., 2001; Rodriguez-Mateos et al., 2009), but with some aggregation of ribosomes as a small single punctate structure located in each conidium cell. Examination of the Rpl25 amino acid sequence predicted two nuclear localisation (NLS) sequences and co-expression of Rpl25-GFP with red fluorescent H1:RFP confirmed that the punctate structure resides in the nucleus of each cell (Supplementary Fig. S1C), likely within the nucleolus. The mitochondria marker, Scad2-GFP, localised to elongated tubular structures evenly distributed in the conidium (Fig. 1D) typical of mitochondrial networks in fungal cells (Aliyu et al., 2019; Kilaru et al., 2017; Suelmann and Fischer, 2000), which, as expected, co-localised with the mitochondrial selective dye; MitoTracker^TM^ Red CM-H2Xros (Invitrogen TM, ThermoFisher Scientific, Waltham, MA, USA) (Supplementary Fig. S1C). The retention signal peptide HDEL (Pelham, 1990) was used for visualisation of the endoplasmic reticulum (ER). Subcellular visualisation of GFP-HDEL fusion in *M. oryzae* revealed that it localised to a reticulate network in the cytoplasm and ring-like peri-nuclear ER structure around the nucleus (Fig. 1E), which co-localised with the ER-specific dye ER-tracker Blue DPX (Supplementary Fig. S1D), consistent with previous observations in *Ustilago maydis* (Wedlich-Soldner et al., 2002), *Zymoseptoria tritici* (Kilaru et al., 2017) and *S. cerevisiae* (Sugiyama and Tanaka, 2019). Localisation of Vac8-GFP (He et al., 2012) was observed in large spherical clusters within conidia (Fig. 1F) consistent with vacuoles previously reported in *M. oryzae* (He et al., 2012) and *U. maydis* (Steinberg and Schuster, 2011). Early Golgi bodies were observed by expression of Grh1-GFP to puncta within conidia (Fig. 1G). A similar localisation was observed for the *trans*-Golgi network (TGN) marker Sec7-GFP (Fig. 1H), consistent with observations in *Aspergillus nidulans* (Pantazopoulou and Peñalva, 2009). The plasma membrane t-SNARE marker, Sso2-GFP, localised to the periphery of conidia with intense expression at septa (Fig. 1I), which co-localised with the lyophilic dye FM4-64 (Kankanala et. al., 2007) consistent with plasma membrane localisation (Supplementary Fig. S1E) as observed in *Z. tritici* (Kilaru et al., 2017). The peroxisome marker Pex6-GFP localised to short filaments in the cytoplasm evenly distributed within conidia (Fig. 1J), as reported previously (Wang et al., 2007). Importantly, we tested the infection ability of all transformants used in this study to ensure that random ectopic integration of individual organelle-tagging vectors did not affect pathogenicity. Conidia from each strain were used in leaf drop infection assays to infect seedlings of susceptible rice cultivar CO39 (Talbot et al., 1993), and were unaffected in their ability to cause rice blast disease (Supplementary Fig. S1F).

**Fig. 1.**
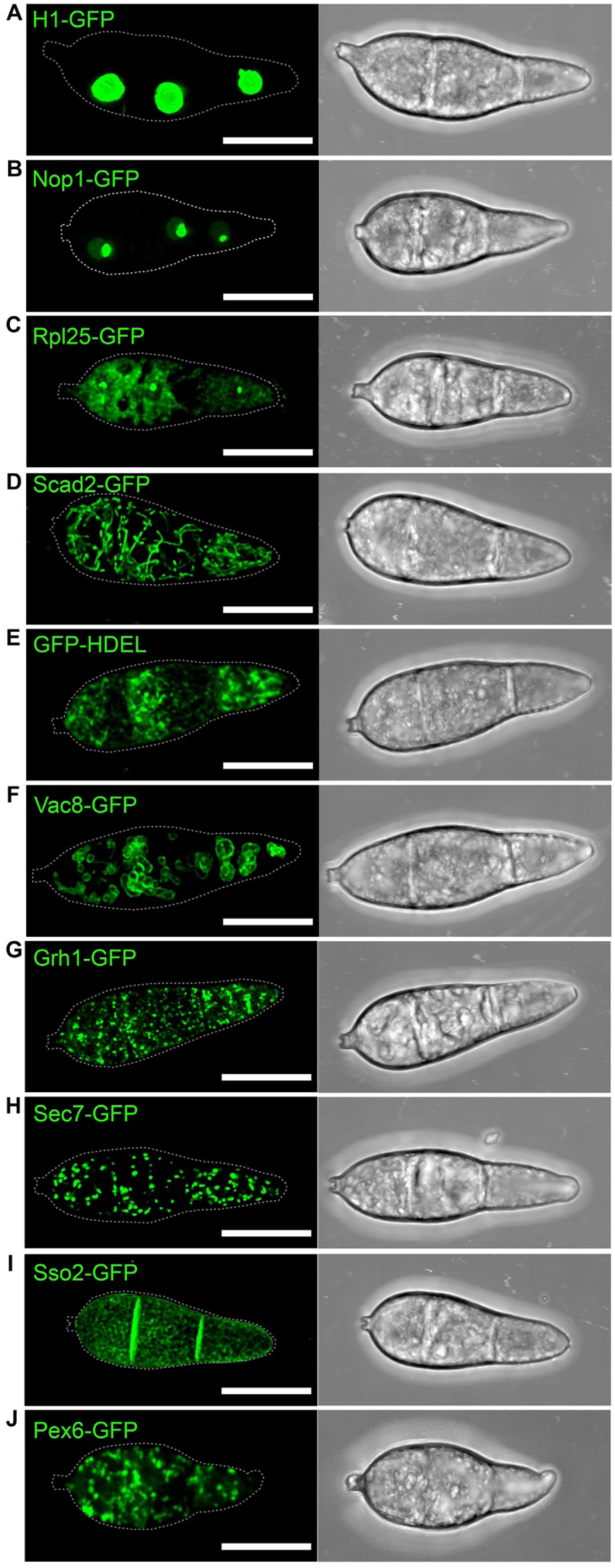
Live cell imaging reveals the sub-cellular localisation of distinct organelle types in conidia of *M. oryzae*. Confocal micrographs of *M. oryzae* Guy11 expressing gene fusions of (**A**) H1-GFP to visualise the nucleus (**B**) Nop1-GFP to visualise nucleoli (**C**) Rpl25-GFP to localise ribosomes. (**D**) Scad2-GFP to visualise mitochondria (**E**) GFP-HDEL to visualise the endoplasmic reticulum, a reticulate structure throughout the conidium. (**F**) Vac8-GFP to visualise vacuoles (**G**) Grh1-GFP to visualise the early Golgi apparatus (**H**) Sec7-GFP to visualise the *trans* Golgi network, (**I**) Sso2-GFP to visualise the plasma membrane (**J**) Pex6-GFP to visualise peroxisomes. Scale bars represent 10 µm.

### Sub-populations of organelles are trafficked to the incipient appressorium

To investigate the fate of each organelle within the three-celled conidium during appressorium development, we performed 4D live-cell imaging and quantitative analysis to determine spatial organisation of each organelle. First, we observed nuclear dynamics using a *M. oryzae* strain expressing H1-GFP (Supplementary Fig. S2A. We observed a single round of mitosis, followed by degradation of all conidial nuclei, consistent with previous reports (Eseola et al., 2021; Saunders et al., 2010a; Veneault-Fourrey et al., 2006a). We then examined nucleolar dynamics using *M. oryzae* Guy11 strains expressing Nop1-GFP (Fig. 2A; Supplementary Video S1). Mitosis occurred by 4 h with a daughter nucleus and nucleolus moving to the incipient appressorium and one to the germinating conidial cell and the majority of germlings containing 4 nuclei. Nuclear degradation then occurred sequentially from the distal non-growing conidium cell to the germinating cell at 8 h (Figure 2A). By 24 h, the conidium was devoid of fluorescence, resulting in a single nucleus and nucleolus being present in the appressorium (Fig. 2A), consistent with the onset of regulated cell death ((Eseola et al., 2021; Saunders et al., 2010a; Shen et al., 2020; Veneault-Fourrey et al., 2006a).

**Fig. 2.**
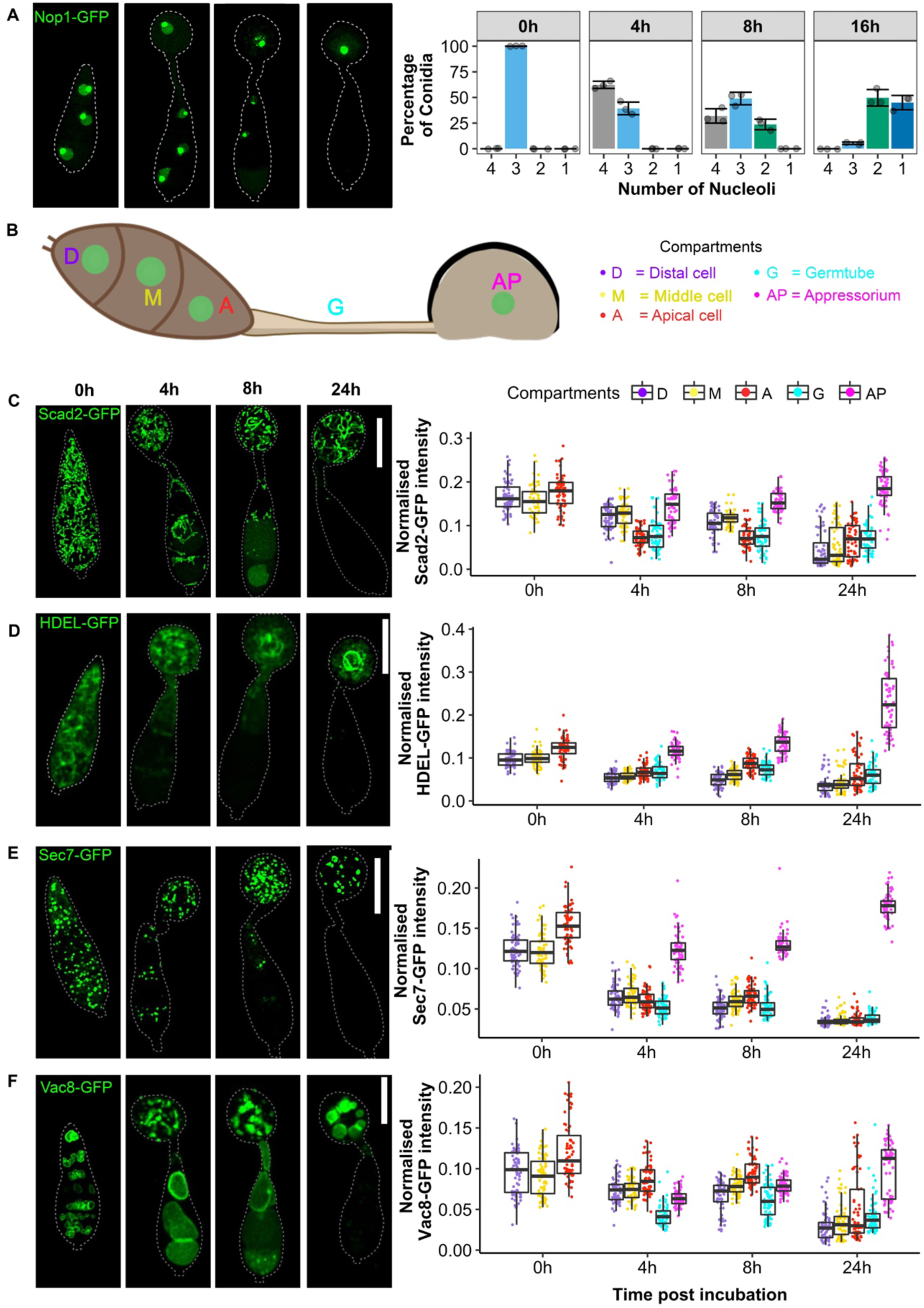
A sub-population of organelles is trafficked to the appressorium during development. **(A)** Representative fluorescence micrographs showing nucleolar dynamics (Supplementary Video S1) and a corresponding bar chart showing nucleolar dynamics during appressorium development of *M. oryzae.* (B) Image illustrating the labelling of the different cellular compartments of *M. oryzae*, which was used to quantify fluorescence intensity. (C) Micrographs of *M. oryzae* conidia expressing mitochondrial marker; Scad2-GFP and the corresponding boxplots to show mitochondria dynamics (Supplementary Video S2) at specific time of appressorium development. (D) Micrographs of *M. oryzae* conidia expressing ER marker; GFP-HDEL and the corresponding boxplots to show ER dynamics (Supplementary Video S3) during appressorium development of *M. oryzae*. (E) Micrographs of *M. oryzae* conidia expressing TGN marker; Sec7-GFP and the corresponding boxplots showing TGN dynamics (Supplementary Video S4) at specific time of appressorium development. (F) Micrographs of *M. oryzae* conidia expressing vacuolar marker; Vac8-GFP and the corresponding boxplots to show the dynamics of vacuoles (Supplementary Video S5) at specific time of appressorium development. The images are maximum intensity projections of Z-stacks taken at 0h, 4h, 8h, and 24h after incubation. The white dashed line outlines the cell boundary drawn with ImageJ; the scale bar represents 10 µm. Boxplots show organelle population within each cell compartment estimated by the mean normalised quantified fluorescence intensity of each organelle marker. The different coloured points indicate each conidium compartment (N = 25 conidia, 3 replicates). The horizontal line of the boxplot is the mean of three replicates and the box represents the quantile, the error bar represents the standard deviation.

We investigated the dynamics of all other organelles during appressorium morphogenesis by performing high-resolution time-lapse imaging using *M. oryzae* strains expressing each individual organelle marker. This revealed that shortly after germination, a polarised germ tube extended from the conidium by 2 h after germination. Mitochondria (Supplementary Video S2), ER (Supplementary Video S3), TGN (Supplementary Video S4), vacuoles (Supplementary Video S5), ribosomes (Supplementary Video S6), early Golgi bodies (Supplementary Video S7) and peroxisomes (Supplementary Video S8) all moved towards the germ tube tip and from 3 h, an influx of organelles from the conidium into the expanding incipient appressorium was consistently observed. The continuous increase of organelles in the incipient appressorium simultaneously led to their decrease in the conidium and by 8 h, the distal cell showed little visible green fluorescence associated with any organelle type (Fig 2C-F; Supplementary Fig. S2C-E, 8 h timepoint) and a decrease in organelle population was also observed in the middle cell (from 9 h). Cell death of the conidium occurred from 12 h by which time organelles were only present in the appressorium.

To evaluate temporal dynamics of organelle trafficking within each conidial cell, we established a quantitative technique to monitor fluorescence intensity in each cellular compartment over time. First, we defined the distal conidium cell (D), middle cell (M), apical cell (A), germ tube (G) and appressorium (AP) as shown in Fig 2B and Supplementary FigS2B. We next carried out time-specific imaging and quantified fluorescence intensity in each compartment. This revealed a continuous increase in mitochondria, ER, TGN, and vacuoles (Fig. 2C-F), ribosomes, early Golgi, and peroxisome movement (Supplementary Fig. S2C-E) to appressoria (AP) over time with a concomitant To model the changes in fluorescence by time and cell region, we created a simple linear regression to plot the abundance of organelles within each compartment using the GFP intensity data from Fig. 2B-F. This model showed a linear decrease in fluorescence intensity of GFP-tagged organelles in compartments D and M during appressorium development (Supplementary Fig. S3). By contrast, fluorescence intensity in the apical conidial cell (A) consistently decreased at a much slower rate with a simultaneous linear increase in organelle abundance in the incipient appressorium (AP) (Fig. S3). All organelles therefore appear to be trafficked to (or are newly synthesised in) appressoria while their dynamics in conidia correspond to the onset of regulated cell death.

Next, we visualised the integrity of the plasma membrane during appressorium development and conidial regulated cell death. For this, we carried out time-specific live-cell imaging using *M. oryzae* expressing the plasma membrane marker, Sso2-GFP, under control of its native promoter. At 0 h, Sso2-GFP localised to both the periphery of the conidium and septa. By 2 h after germination, the germ tube extending from the germinating conidium displayed Sso2-GFP signal at its periphery, which was weak at the germ tube tip (Supplementary Fig. S4A). From 4 h, following formation of the incipient appressorium, Sso2-GFP was limited to the appressorium collar, and no visible fluorescence was detected at the appressorium cell cortex (Figure S4A, 4 h). We reasoned that this might be due to the presence of melanin at the appressorium preventing effective visualisation of the plasma membrane. At 8h, we observed disintegration of the conidial cell membrane, starting with the distal conidia cell at 8 h, middle cell by 12 h, and apical and germ tube by 24 h (Figure S4A). Complete disintegration of the conidial plasma membrane occurred by 24 h, indicating completion of conidial cell death (Veneault-Fourrey et al., 2006a). To investigate whether masking of plasma membrane fluorescence in the appressorium was associated with the presence of the melanin layer in the appressorium cell wall, we expressed Sso2-GFP in a melanin-deficient mutant *alb1* (Foster et al., 2018). This revealed Sso2-GFP at the appressorium cortex at 24h (Figure S4B), confirming that melanisation masked expression of Sso2-GFP in appressoria of the wildtype strain, Guy11. We conclude that conidial cell death occurs sequentially from the distal cell to the apical cell of the conidium and is completed by 24h after germination.

### Organelles are actively transported to the appressorium

4D live cell imaging provided evidence that organelles are trafficked to the appressorium of *M. oryzae,* but it was not possible to determine whether organelles in the appressorium originated in the conidium, were newly synthesized, or a combination of both. We therefore used fluorescence recovery after photobleaching (FRAP) (Jacobson et al., 1991) to monitor fluorescence recovery at the photobleached germ tube tip during appressorium formation. We photobleached the swelling tip of the germ tube to monitor the rate of fluorescence recovery. Diffuse movement of intact mitochondria (Fig. 3A; Supplementary Video S9), the TGN (Fig. 3B; Supplementary Video S10), and ER (Fig. 3C; Supplementary Video S11) were observed to move to the photobleached incipient appressorium. We observed a 56% recovery of Scad2-GFP fluorescence in the germ tube after 35 min, a 66% recovery of Sec7-GFP fluorescence after 35 min and a 99% recovery of GFP-HDEL after 15 min. GFP-HDEL appeared to recover most rapidly, suggesting the ER is highly dynamic. We conclude that active transport of organelles occurs during appressorium development (Figure 3A-C).

**Figure 3.**
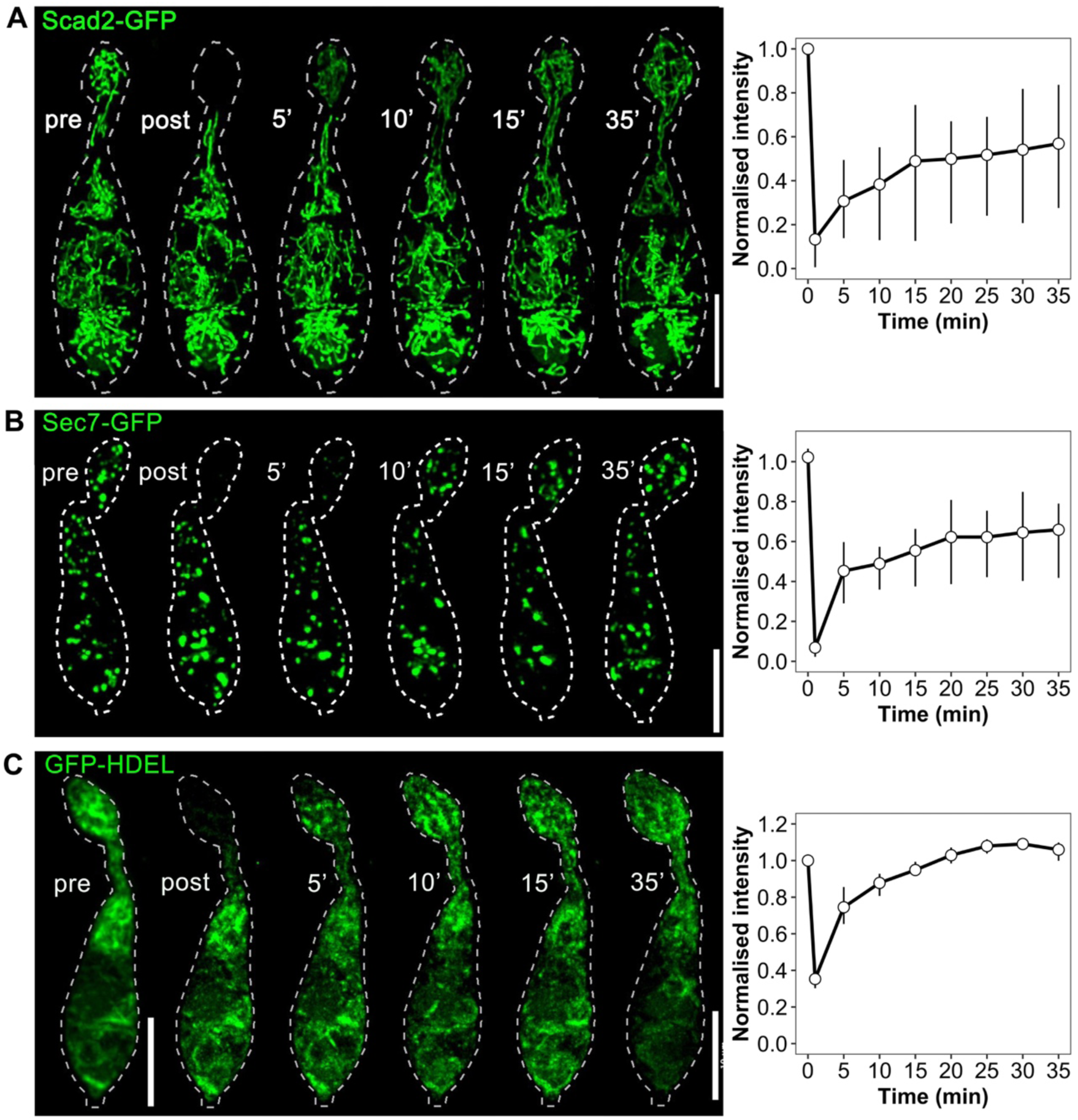
FRAP analysis shows active transport of GFP into photobleached germ tube tips of *M. oryzae*. Representative still images of (A) Scad2-GFP (Supplementary Video S9), (B) Sec7-GFP (Supplementary Video S10) and (C) GFP-HDEL (Supplementary Video S11) recovery in the swollen germ tube tips after photobleaching. Conidia were incubated on a hydrophobic coverslip for 2h before conducting FRAP analysis. Micrographs are maximum intensity projections of *Z*-stack series. Recovery time is displayed in minutes, and the scale bar represents 10 µm. The graph on the left shows the average fluorescence recovery curves for each fluorescence marker. Results are presented as mean and standard error of the mean calculated from 3 biological replicates.

To define the likely mechanism of organelle trafficking, we exposed developing appressoria to benomyl and latrunculin-A, which depolymerise microtubules and F-actin filaments, respectively. We tested concentrations of benomyl (10 µg mL^-1^) and latrunculin-A (10 µg mL^-1^) previously shown to be sufficient to depolymerise the microtubule network and F-actin organisation of *M. oryzae* strains expressing β-tubulin-GFP (Saunders et al., 2010a) and TpmA-GFP (Saunders et al., 2010b), respectively. We observed depolymerisation of microtubules when conidia expressing β-tubulin-GFP were exposed to benomyl at 4 h (Figure S5A), whereas latrunculin-A and DMSO, a solvent control, did not depolymerise microtubules (Figure S5A). In contrast, we observed mislocalisation of the F-actin disc, which normally forms at the appressorium pore (Dagdas et al., 2012), when conidia expressing TpmA-GFP were exposed to either benomyl or latrunculin-A at 4 h (Figure S5B). By contrast, the F-actin disc was organised at the centre of the appressorium in the DMSO-treated sample. Next, we examined mitochondrial transport into the appressorium by treating conidia of *M. oryzae* strain expressing Scad2-GFP with either inhibitory drug at 4 h. We observed that normal elongated mitochondrial morphology was severely fragmented upon exposure to benomyl (Figure S5C) and by 24 h, benomyl-treated samples also displayed non-melanized appressoria, suggesting that disruption of microtubules prevented transport of melanin precursors to the appressorium or had wider effects on development. Similarly, we tested whether actin filaments are required for short-range mitochondrial transport by exposing conidia of Scad2-GFP strain to 10 µM latrunculin-A at 24 h. Disruption of actin filaments had less of an effect on mitochondria morphology initially, but a reduced population was observed in the appressorium by 24 h with some remaining in the germ tube and conidium (Figure S5D). To estimate the effect of cytoskeleton disruption, we compared the sub-population of mitochondria in appressoria at 24h which showed a significant reduction (*P* < 0.001) in the mitochondrial population in both benomyl-treated (Figure S5E) and latrunculin-A treated (Figure S5F) samples compared to a DMSO control sample (Figure 5.5C and F). Taken together, we conclude that the microtubule and actin cytoskeleton is involved in movement of mitochondria to the appressorium in *M. oryzae,* consistent with previous reports of motor-dependent organelle trafficking in fungi (Lin et al, 2016).

### Spatial diffusion of photoactivatable GFP demonstrates that the germinated three-celled conidium is a contiguous open system

To investigate whether the three single cells that comprise a conidium in *M. oryzae* are interconnected by open septal pores, we carried out conditional photoactivation of individual conidial cells to monitor cytoplasmic movement between them. Cells of filamentous fungi contain a septum, which defines each cell but still enables intercellular communication via cytoplasm and organelle exchange through septal pores (Abadeh and Lew, 2013). It has been previously reported that Woronin bodies (also known as occlusion bodies) in *M. oryzae* do not block conidium septal pores during early stages of appressorium development (Soundararajan et al., 2004). We therefore reasoned that if septal pores in the conidium are open, immediate trafficking of a reporter marker should be observed throughout the three-celled conidium. To test this idea, we expressed cytosolic photoactivatable GFP, paGFP (Patterson and Lippincott-Schwartz, 2002) under a constitutive promoter. We then differentially pulsed the distal, middle and apical cell, respectively with a 405 nm laser (excluding the septum) to activate the reporter. The photoactivated cell compartment instantly fluoresced and enabled spatial monitoring of paGFP by time-lapse imaging. After pulsing the distal cell (Figure 4A, Supplementary Video S12), photoactivated paGFP was immediately observed in the germ tube and within 5 min, paGFP was visible throughout the three-celled spore. A similar observation occurred when the middle (Figure 4B, Supplementary Video S13) and apical (Figure 4C, Supplementary Video S14) cells were individually pulsed. These observations provide strong evidence that the three-celled conidium of *M. oryzae* is initially an open system connected by septal pores.

**Figure 4.**
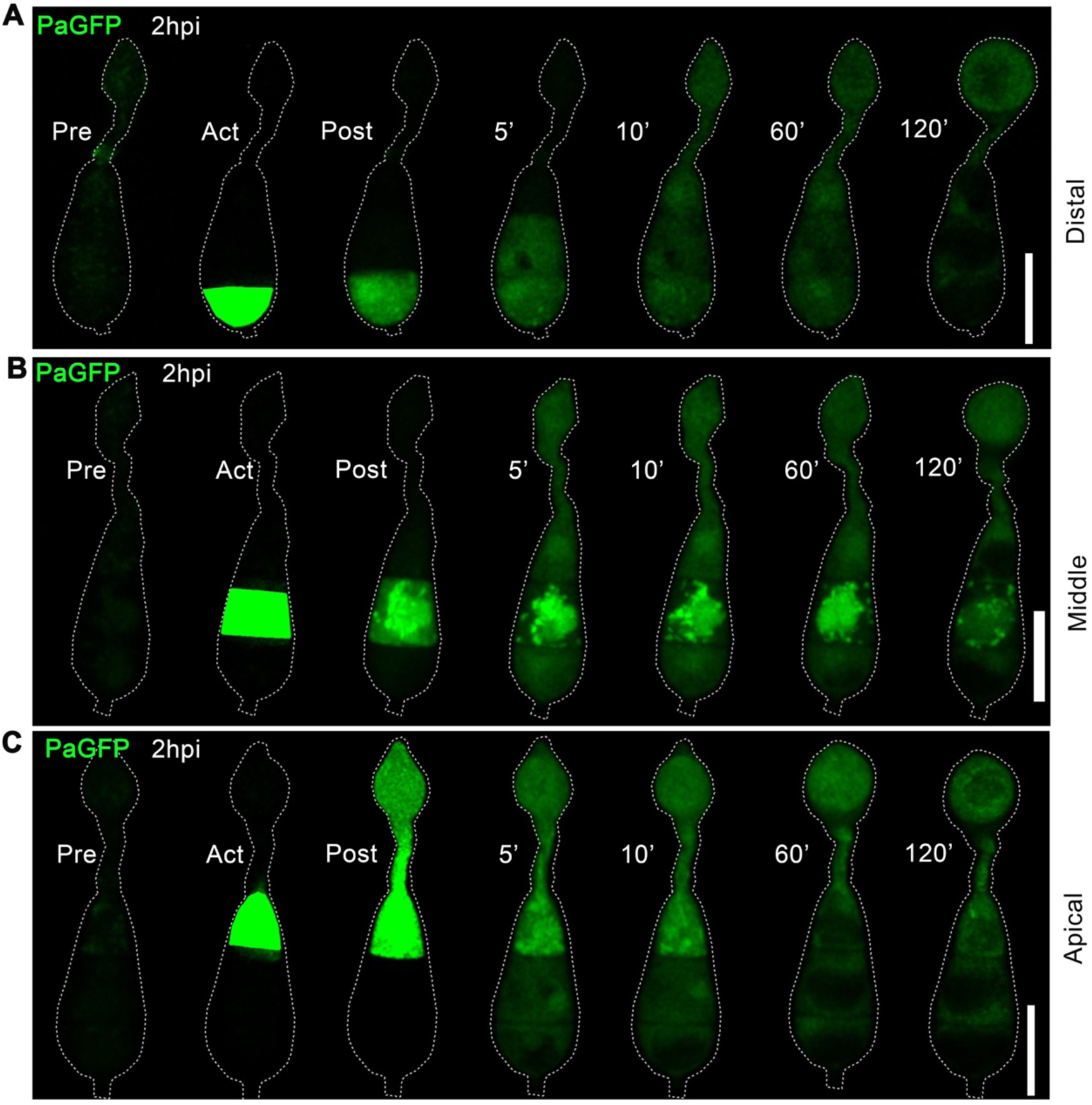
Photoactivation of paGFP confirms that the three celled conidium is an open system connected by open septal pores. Conidia expressing paGFP were incubated on a hydrophobic coverslip for 2 h before photoactivation. Before photoactivation, paGFP expression was not visible in (A) distal cell; (Supplementary Video S12), (B) middle cell; (Supplementary Video S13) and (C) apical cell; (Supplementary Video S14). After a short pulse with 405-nm laser, fluorescent cytoplasm appeared to traffic into adjacent cells. Micrographs are maximum intensity projections of *Z-*stack series captured at 5 min intervals for 2 h. The dotted line denotes the cell outline and was performed in ImageJ. The displayed stilled frames are maximum intensity projections of the *Z*-stack series. Time is displayed in minutes, and the scale bar represents 10 µm.

**Figure 5.**
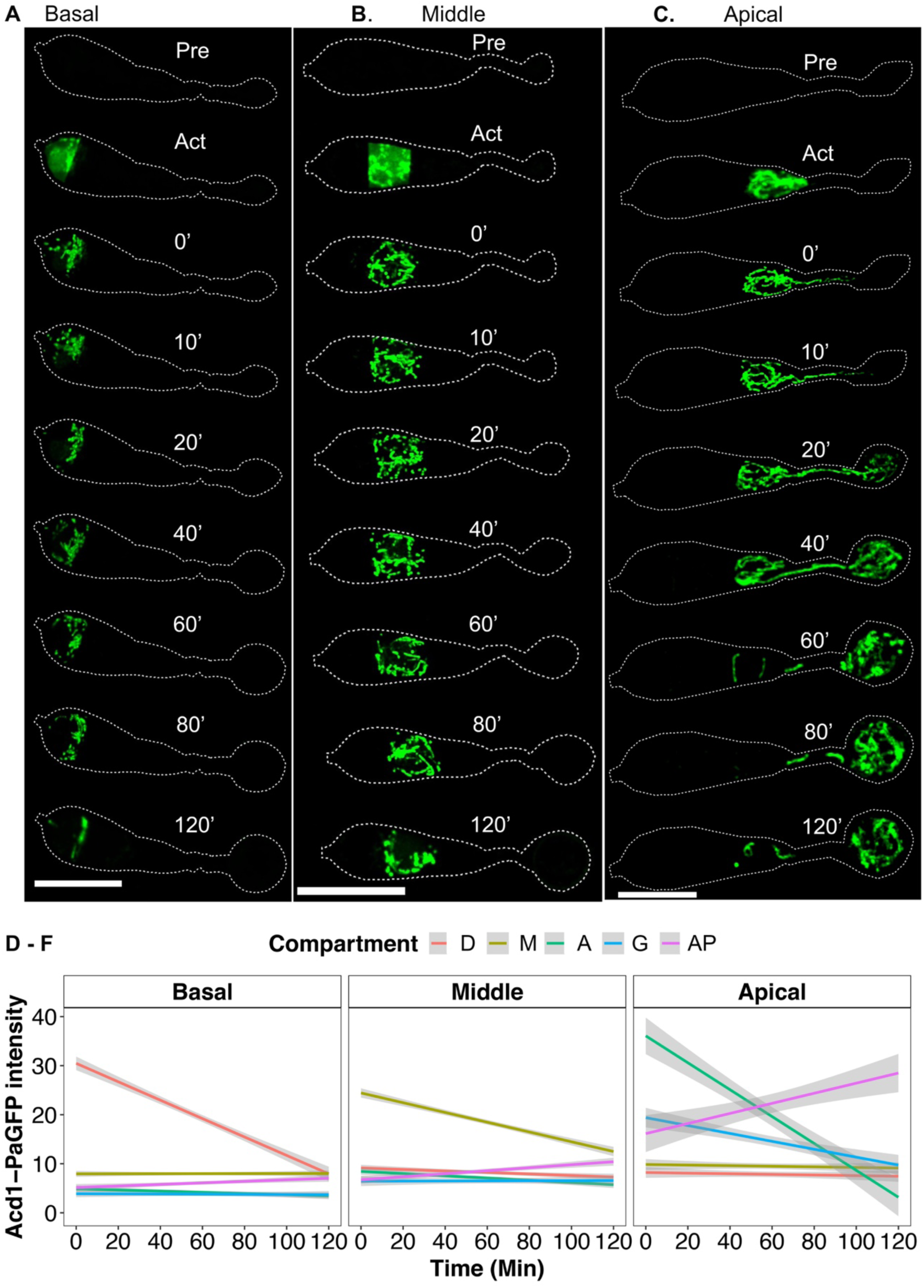
Photoactivation reveals that mitochondrial movement into the appressorium occurs only from the growing apical cell of the conidium. Conidia expressing Scad2-paGFP were incubated on a hydrophobic coverslip for 2 hours before photoactivation. Prior to photoactivation, mitochondria were not visible in conidia. After a short pulse with a 405-nm laser for 5 sec, fluorescent mitochondria in (A) distal (Supplementary Video S17) and (B) middle (Supplementary Video S18) showed no transport into adjacent compartments. (C) Fluorescent mitochondria in the apical compartment (Supplementary Video S19) showed trafficking through the germ tube to the appressorium. Micrographs are maximum intensity projections of *Z-*stack series captured at 5 min intervals for 2 h. The dotted line outlines the cell and performed in ImageJ. Time is displayed in minutes, and the scale bar represents 10 µm. (D-F) Simple linear regressions of fluorescence intensity over time in given cell regions show the dynamic distribution and fate of photoactivated mitochondria in the distal, middle, and apical conidia cells during 2 h. The mean is calculated from three biological replicates. The shaded area of the graph represents the standard error.

Following completion of mitosis, an asymmetric cytokinesis event results in a septum that delineates the developing appressorium (Saunders et al., 2010b), which becomes competent to generate turgor. To investigate whether the appressorium is sealed from the collapsing conidium after this cell division event, we examined movement of cytosolic paGFP. We first pulsed paGFP in the conidium at 5h to monitor cytoplasmic streaming into the incipient appressorium. We observed that cytosolic paGFP diffused into adjacent conidium compartments and the appressorium (Fig. S6A; (Supplementary Video S15). However, when we pulsed the apical conidium at 6h, cytosolic paGFP was restricted to the appressorium collar (Supplementary Fig. S6B; (Supplementary Video S16). This result suggests that cytoplasmic diffusion into the appressorium is prevented after sealing of the appressorium collar by completion of the septation event (Saunders et al., 2010b). We conclude that the germinating conidium is an open system that facilitates trafficking from all cells to the appressorium until mitotic exit and appressorium cytokinesis, which are complete by 6 h after conidial germination.

### Organelle trafficking is not uniformly regulated from each conidial cell to the appressorium

To determine whether the three-celled conidium of *M. oryzae* uniformly contributes organelles to the developing appressorium, we carried out live-cell imaging using a *M. oryzae* strain expressing the mitochondria marker Scad2 fused to paGFP to monitor organelle dynamics during appressorium morphogenesis. Selective excitation of Scad2-paGFP in the distal (Supplementary Video S17) Figure 5A) and middle (Supplementary Video S18) Figure 5B) cells of the conidium did not result in any mitochondrial movement to neighbouring compartments. However, the Scad2-paGFP signal subsequently decreased, consistent with organelle breakdown. By contrast, when we monitored Scad2-paGFP in the apical conidial cell, mitochondria instantly moved through the germ tube to the appressorium (Supplementary Video S19); Figure 5C). To quantify the behaviour, we plotted fluorescence intensity of Scad2-paGFP within each compartment and used a linear graph to visualize the rate of trafficking and/or simultaneous degradation within each conidial cell. A continuous linear degradation of Scad2-paGFP was observed in distal and middle cells of the conidium (Figure 5D-E) following paGFP activation. By contrast, a continuous linear increase occurred in the appressorium was observed following apical cell Scad2-paGFP activation (Figure 5F). Taken together, we conclude that the apical cell is the single source of intact mitochondria that are actively trafficked to the appressorium, while organelle breakdown occurs in the distal and middle conidial cells.

### Autophagy occurs sequentially in non-germinating cells of the conidium

Next, we set out to investigate whether autophagy is necessary for organelle breakdown in the distal and middle cells of the conidium. For this, we expressed Scad2-paGFP in a *Δatg8* null mutant to generate a *Δatg8:*Scad2-paGFP strain unable to undergo autophagy-mediated conidial cell death (Kershaw and Talbot, 2009; Veneault-Fourrey et al., 2006b). Photoactivation of *Δatg8:*Scad2-paGFP in the distal cell revealed no trafficking of mitochondria to neighbouring cell and no breakdown after 2 h ((Supplementary Video S20); Fig. 6A). Similarly, photoactivation of Scad2-paGFP in the middle cell of the *Δatg8* mutant also resulted in signal restricted to the individual cell with no visible organelle degradation (Supplementary Video S21; Fig. 6B). However, photoactivation of Scad2-paGFP in the apical cell resulted in migration of fluorescent mitochondria to the incipient appressorium (Supplementary Video S22); Fig. 6C). We conclude that autophagy is necessary for organelle breakdown in the distal and middle conidial cells, but that organelle trafficking to the appressorium from the apical cell occurs prior to autophagy being initiated in this compartment. This suggests that spatio-temporal regulation of autophagy occurs in the conidium such that distal and middle cells undergo autophagy and cell death ahead of the germinating apical cell.

**Figure 6.**
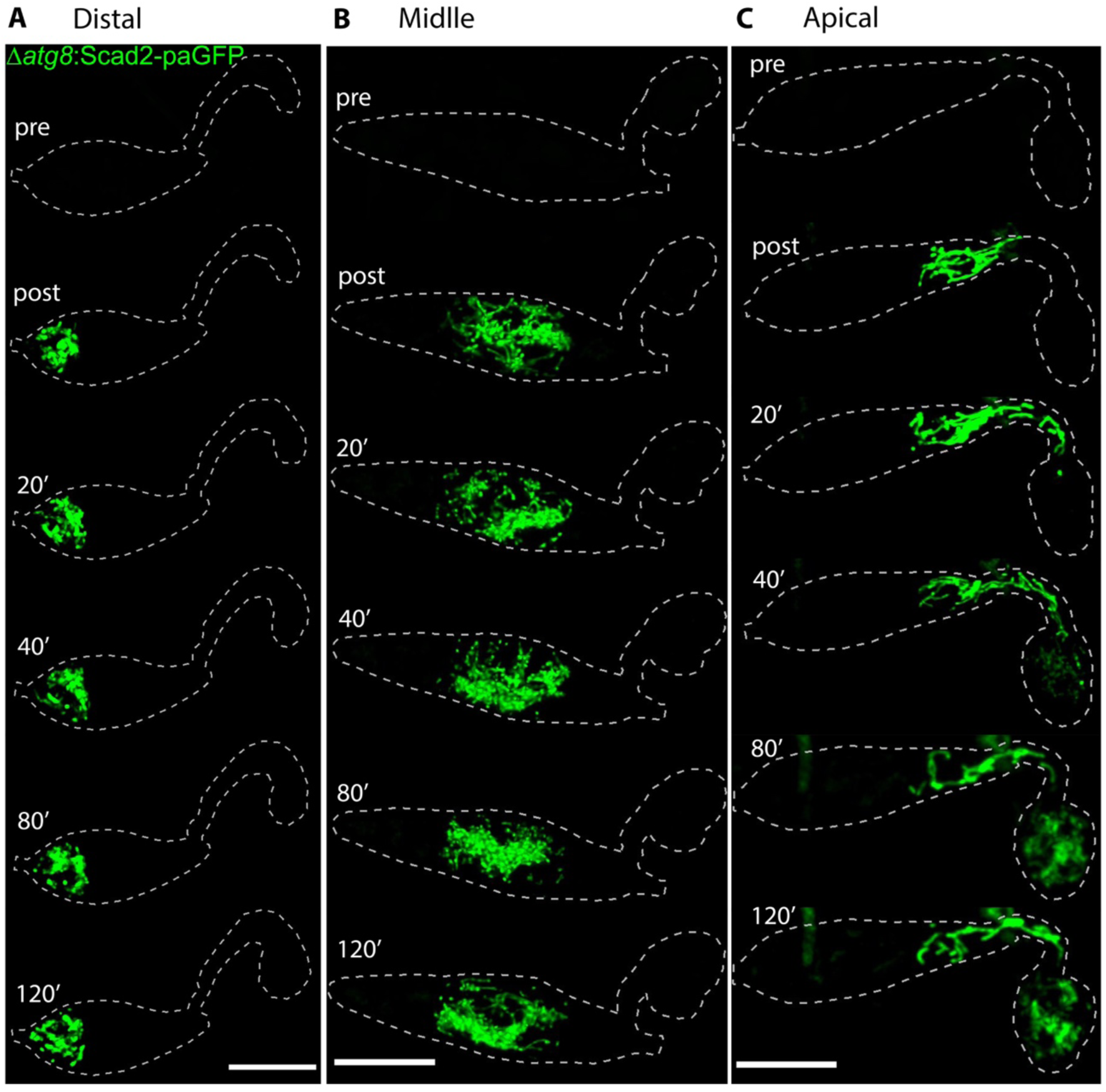
Autophagy is not necessary for trafficking towards the appressorium from apical cell, but it is necessary for organelle degradation from middle and distal cells. Representative micrographs of *M. oryzae Δatg8* null mutant conidia expressing Scad2-paGFP were harvested and incubated on a hydrophobic coverslip for 4h. Before photoactivation, mitochondria were not visible in the conidia. After a short pulse with a 405-nm laser, fluorescent mitochondria in the (A) distal and (B) middle cells showed no transport into other compartments and were not degraded. (C) Fluorescent mitochondria in the apical conidium cell traffic into the appressorium. Micrographs are maximum intensity projections of *Z*-stack series captured at 20 min intervals for 2 h. Micrographs are till frames taken from (Supplementary Video S20), (Supplementary Video S21) and (Supplementary Video S22) respectively. The dotted line outlines the cell and was performed in ImageJ. Time is displayed in minutes, and the scale bar represents 10 µm.

To determine whether the same observations occur with distinct organelle types, we expressed H1-GFP, Nop1-GFP, GFP-HDEL, Rpl25-GFP, Sec7-GFP, Grh-GFP, Pex6-GFP and Sso2-GFP in the *Δatg8* mutant background (Supplementary Fig. S7). Organelle breakdown was impaired in all cases, resulting in distorted and aberrant organelle morphology in conidial cells (Supplementary Fig. S7). Appressorium development occurred over an 8 h period in the *Δatg8* mutant background and by 24 h nuclei, nucleoli, mitochondria, ER ribosomes, Early Golgi, TGN, and peroxisomes (Supplementary Fig S7A-H) showed distorted morphology and organisation in conidia. Plasma membrane dissolution in conidial cells revealed by Sso2-GFP fluorescence was also impaired (Supplementary Fig S7I), consistent with the lack of cell death. Taken together, these observations suggests that autophagy is important for organelle degradation but does not affect organelle inheritance from the apical conidial cell to the appressorium.

### Organelle inheritance by the appressorium is independent of cell cycle control in *M. oryzae*

In *M. oryzae*, three cell cycle checkpoints regulate development and function of the appressorium (Osés-Ruiz et al., 2017; Saunders et al., 2010a). We reasoned that organelle trafficking from the apical cell towards the incipient appressorium might be cell cycle-regulated and we therefore expressed Scad2-paGFP in conditional mutants of *M. oryzae* impaired at specific phases of the cell cycle. First, we examined whether the S-phase checkpoint controlled organelle trafficking by using a temperature-sensitive mutant in the B-type cyclin gene *cyc1^nimE10^*(Osés-Ruiz et al., 2017). We incubated conidia of *cyc1^nimE10^*:Scad2- paGFP in a pre-heated incubator at a semi-restrictive temperature of 30°C to arrest the mutant at S-phase before being imaged in a pre-heated microscope chamber for another 3 h. Conidia of *cyc1^nimE10^*:Scad2-paGFP produced a polarised germ tube at 2 h and each individual conidial cell was then pulsed with high-intensity light using a 405-nm laser diode to monitor photoactivated mitochondria. We observed Scad2-paGFP in the apical conidia cell still actively trafficked to the germ tube tip (Figure 7A), even though this was unable to differentiate into an appressorium. However, photoactivated mitochondria were not trafficked from the middle (Figure S8A) and distal (Figure S8B) cells. These observations confirm that entry to S-phase is essential for appressorium development (Saunders et al., 2010), but that organelle trafficking occurs independently from the germinating conidial cell.

**Figure 7.**
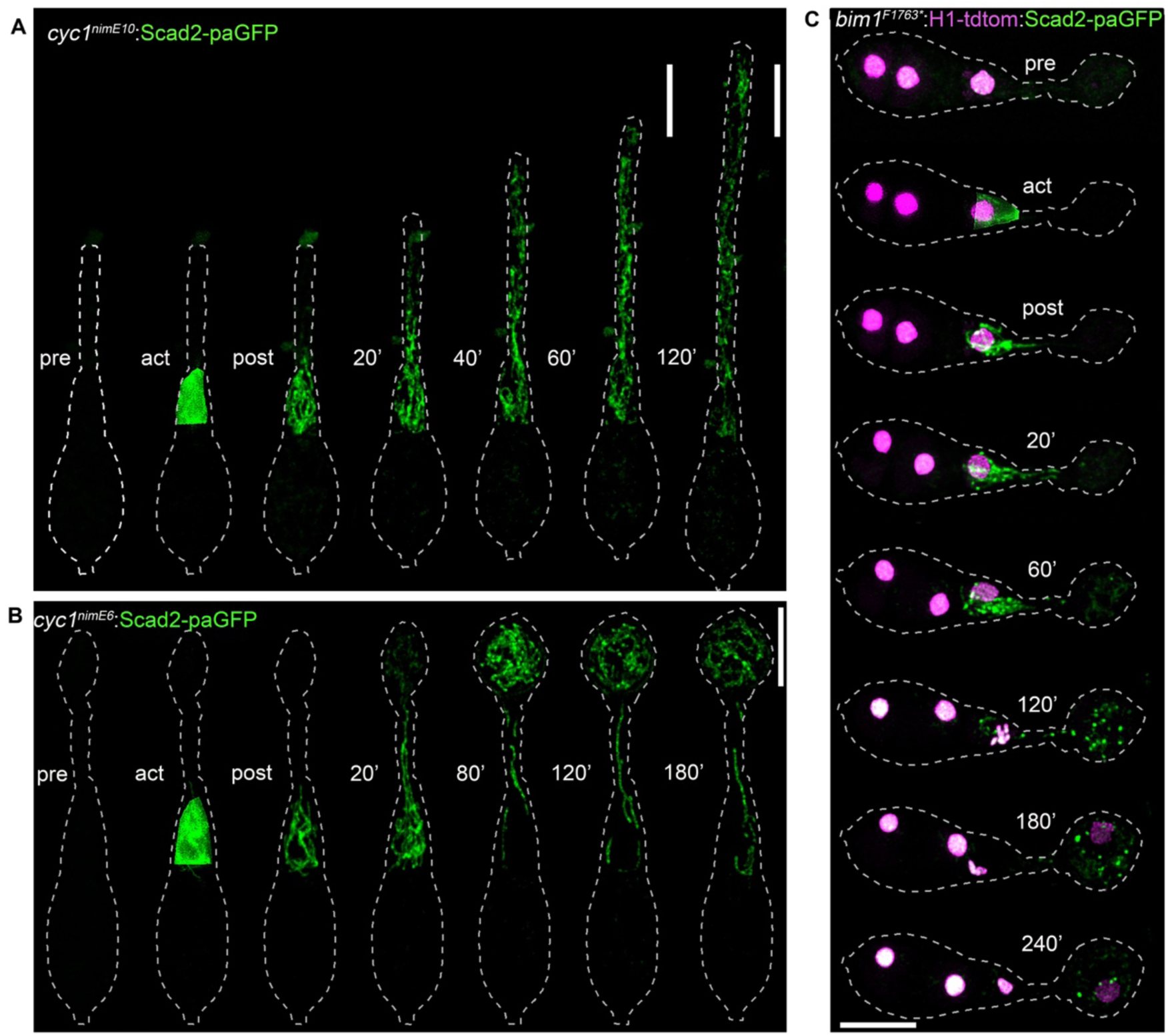
Organelle trafficking is not regulated by cell cycle checkpoints in M. oryzae. (A) Micrographs of conidia of conditional S-phase mutant expressing *cyc1^nimE10^*:Scad2-paGFP, after photoactivation of the apical conidium cell, fluorescent mitochondria trafficked towards the elongating germ tube, and the germ tube failed to differentiate into an appressorium. (B) Micrographs of conidia of conditional G2-phase mutant expressing *cyc1^nimE6^*:Scad2-paGFP, after photoactivation of the apical conidium cell, fluorescent mitochondria trafficked into the incipient appressorium (C) Micrograph of conidium of conditional mitotic mutant photoactivated in the apical cell 3h post-incubation. After photoactivation, fluorescent mitochondria in the apical cell moved through the germ tube into the incipient appressorium. The conidia of the coniditionl mutants were incubated on hydrophobic coverslips at 30°C before photoactivation and live cell imaging in a preheated microscope chamber. The dotted line outlines the cell and was performed in ImageJ. Time is displayed in minutes post activation, and the scale bar represents 10 µm.

Next, we examined G2-M phase by expressing Scad2-paGFP in the conditional mutant *cyc1^nimE6^* (Osés-Ruiz et al., 2017). Conidia of *M. oryzae* strain *cyc1^nimE6^*:Scad2-paGFP were incubated for 2h at semi-permissive 30°C as described above. At 2 h, a germ tube emerged from the conidium, and the germ tube appeared slightly swollen at the tip (Fig. 7B) indicating hooking and initiation of appressorium formation, consistent with previous reports (Osés-Ruiz et al., 2017). Following Scad2-GFP photoactivation of the apical conidium cell, we observed a sub-population of fluorescent mitochondria moved into the incipient appressorium (Fig. 7B) which continued to develop but did not melanise. Similarly, Scad2-GFP photoactivation in the middle (Supplementary Fig. S8C) and distal (Supplementary Fig. S8D) cells resulted in mitochondria remaining in the same compartment, even after 3h of photoactivation. We conclude that G2-arrest, which impairs appressorium maturation, does not prevent organelle transport to the developing appressorium. Finally, we examined whether a mitotic checkpoint regulates organelle trafficking during appressorium development. We examined the spatial dynamics of mitochondria in an *M. oryzae* conditional mitotic mutant expressing a red fluorescent nuclear marker *bim1^1763^*\*:H1-tdtomato (Saunders et al., 2010b). After 3h incubation at semi-permissive 30°C, conidia of the *bim1^1763*^*:H1-tdtom:Scad2-paGFP strain formed hooked germ tubes that later differentiated into appressoria (Fig. 7C), but which were unable to re-polarise and cause infection and were impaired in autophagic cell death (Saunders et al., 2010a). Following Scad2-GFP photoactivation in the apical cell, a sub-population of fluorescent mitochondria migrated to the incipient appressorium (Fig. 7C). However, Scad2-GFP photoactivation in the middle (Supplementary Fig. S8E) and distal cells (Supplementary Fig. S8F), did not result in mitochondrial movement to adjacent cells and conidial contents were not degraded by autophagy. We conclude that organelle trafficking to the developing appressorium is independent of conidial mitosis. Taken together, these experiments reveal that while appressorium morphogenesis and autophagy are cell cycle controlled, organelle trafficking is an independent process.

### Autophagy is not required for *de novo* organelle biogenesis

Having established that mitochondria are trafficked to the appressorium from the apical cell, we set out to determine whether organelle biogenesis occurs within the infection cell prior to plant infection or results solely from trafficking from the apical conidial cell. To do this, we carried out 3D volumetric analysis of mitochondria using Imaris (Bitplane) by manually outlining each organelle in three dimensions (Supplementary Video S23); Fig. 8A) and color-coding each compartment of the three-celled conidium (Fig. 8B). At 0h, the distal, middle, and apical conidia cells contained an average volume of 19.32 µm^3^, 21.04 µm^3^ and 12.20 µm^3^ of mitochondria, respectively (Fig. 8C). After incipient appressorium formation at 4h, we recorded an average of 18.91 µm^3^ mitochondrial volume in the appressorium. Throughout each developmental stage, the mitochondrial volume in the appressorium increased while simultaneously decreasing in conidia. By 24h, all conidia cells had collapsed, leaving a 28.95 µm^3^ volume of mitochondria in the appressorium (Fig. 8C). The volume of mitochondria in the appressorium was therefore greater than the starting average mitochondrial volume in the apical conidial cell at 0h, suggesting that by 24h *de novo* mitochondrial biogenesis occurs during appressorium maturation in *M. oryzae.* To determine whether conidial autophagy is necessary for organelle biogenesis in appressoria, we measured the volume of mitochondria in an *Δatg8* mutant. At 0h, the distal, middle, and apical cells contained an average volume of 43.7 µm^3^, 56.4 µm^3^ and 31.2 µm^3^, respectively (Supplementary Fig. S9). Mitochondrial volume in the apical cell decreased over time due to transport towards the appressorium. However, mitochondrial volume in the distal and middle cells remained unchanged throughout appressorium development in the *Δatg8* mutant consistent with impaired autophagy (Fig. S9). Mitochondrial volume in the appressorium still exceeded the volume in the apical cell by 24h (66.9 µm^3^). This analysis suggests that organelles undergo biogenesis during transportation to appressorium of *M. oryzae* and that this occurs independently of autophagy.

**Figure 8.**
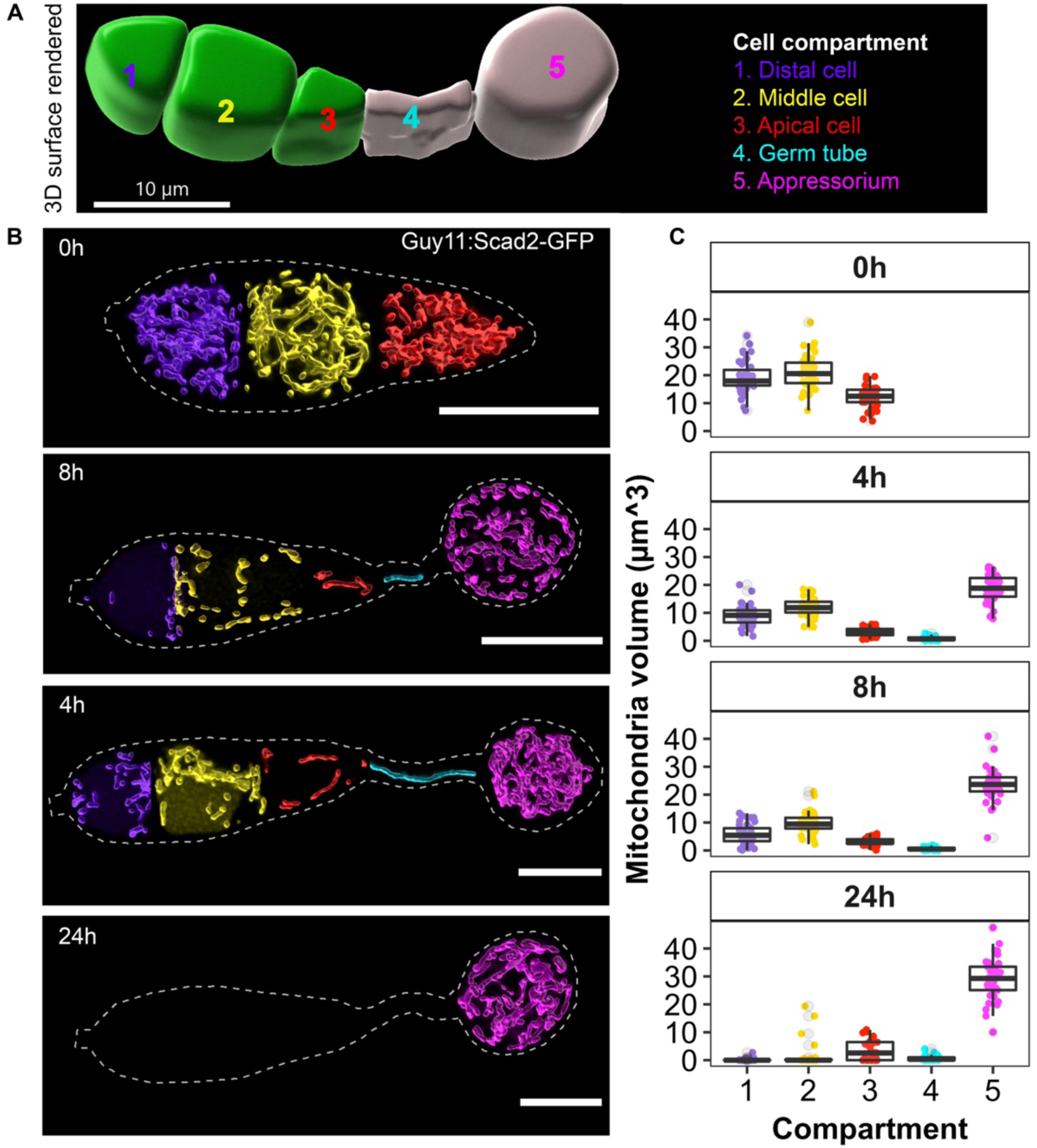
Volumetric analysis of intact mitochondrial during appressorium development of *M. oryzae*. (A) 3D rendered surface of a germinated *M. oryzae* conidium to show how mitochondria volume was quantified within each numbered conidium cell (Supplementary Video S23). (B) Micrographs showing the distribution of intact mitochondria within conidia of the wildtype strain Guy11 during appressorium development at a specific time. The scale bar represents 10 µm. Image rendering was performed in Imaris. (C) Boxplot showing the mean volume of intact mitochondria within conidia of wildtype strain Guy11 of *M. oryzae* during appressorium development at 0h, 4h, 8h, and 24h. The different colours in the plot represent each conidium cell (N= 50 conidia).

### Gene fusion of photoconvertible protein reveals the precise timing of organelle biogenesis in *M. oryzae*

To determine the timing of *de novo* organelle synthesis, we generated a fusion of Scad2 with the photoconvertible protein; mEos3.2, a UV-inducible fluorochrome that changes fluorescence from green to red when pulsed with a 405 nm laser (Wiedenmann et al., 2004). We used live-cell imaging to monitor the photoconverted red and newly synthesised green fluorescence intensity dynamics from 2 h at the onset of germ tube swelling. During incipient appressorium development, photoconverted Scad2-mEos3.2 fluorescence migrated from the conidium to the incipient appressorium (Supplementary Video S24); Fig. 9). However, at 07h:02min:24sec, while the distal and middle compartments were still occupied by photoconverted Scad2-mEos3.2 fluorescence (red mitochondria), a green fluorescence signal began to appear in the germ tube and appressorium (Figure 9, indicated by a green asterisk). Shortly afterwards the intensity of red fluorescence signal in the conidium decreased, at the time of autophagic degradation of converted mitochondria in the middle and distal cells. From 07h:36min (Supplementary Video S24), the mitochondrial population in the apical and appressorium compartments consisted of a combination of green and red fluorescence mitochondria (Fig. 9, indicated by white coloured mitochondria). A new population of mitochondria expressing green fluorescence in the appressorium corresponded with the timing of appressorium melanisation at 8h, consistent with rapid *de novo* biogenesis of mitochondria. After 9h (Fig. 9, (Supplementary Video S24, 8h:56min), the distal cell collapses, followed by the middle cell (13h:34min:24sec). These observations provide further evidence that distal and middle conidial cells are programmed for autophagic degradation, and organelle trafficking and *de novo* mitochondrial synthesis occurs only in the germinating conidium cell and appressorium at the onset of appressorium melanisation.

**Figure 9.**
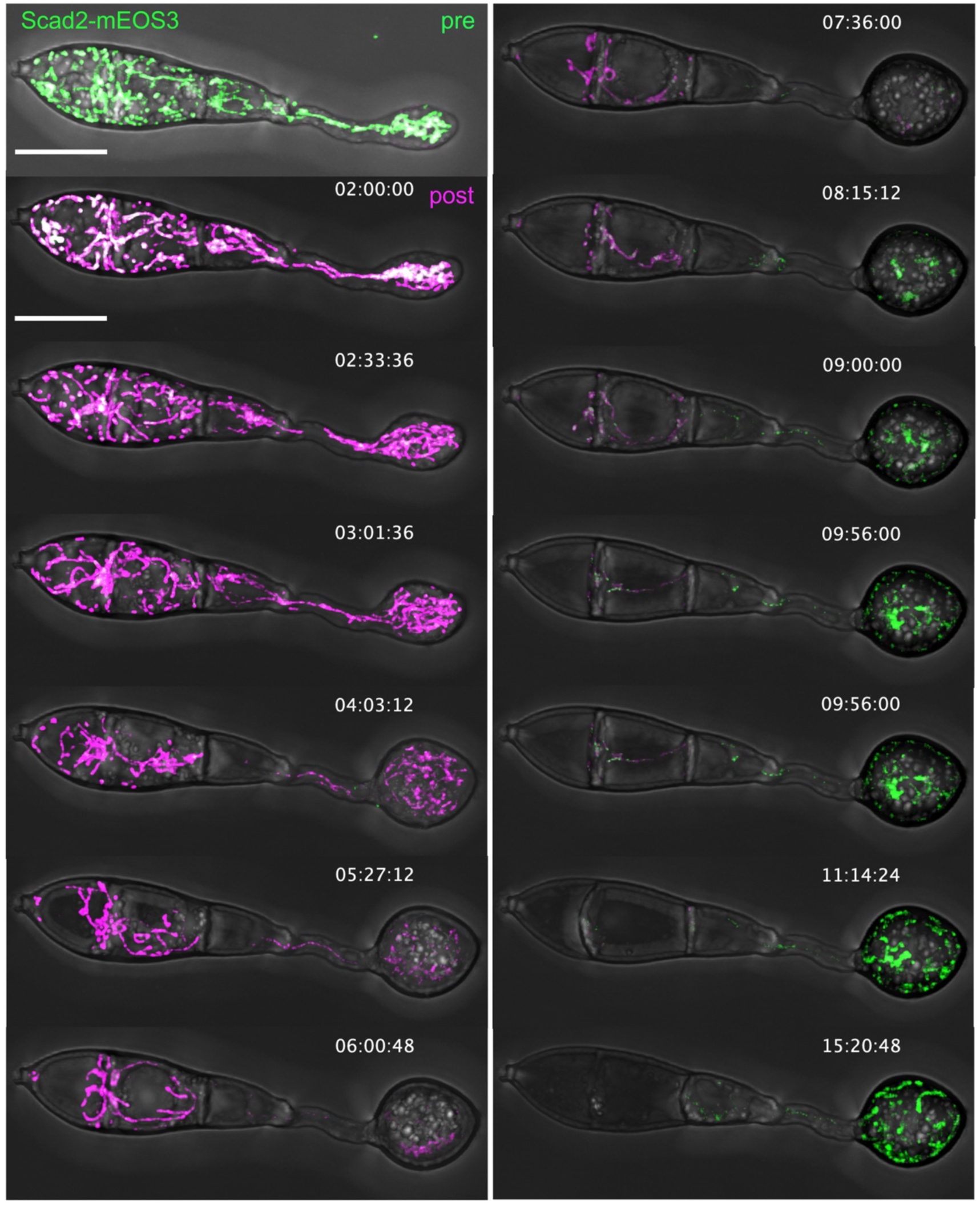
Cellular localisation of a photoconvertible mitochondria marker during appressorium development of *M. oryzae*. Conidia expressing Scad2-mEOS3.2 were harvested and incubated on a hydrophobic coverslip for 2 h before photoconversion. Conversion was achieved by pulsing the conidium with a 405-nm laser diode at 40 % of full power for 5 sec. Green fluorescent images were excited at 488-nm and red fluorescent images were excited with 561-nm. The asterisks indicated the timing of *de novo* biogenesis of mitochondria. Micrographs are three-dimensional projections of *Z-*stack series captured at 5 min intervals, taken from (Supplementary Video S24). Time is displayed in hour:minute:second, and the scale bar represents 10 µm. Time is displayed in hour:minute:second, and the scale bar represents 10 µm..

### Conidial cell germination is necessary and sufficient for organelle trafficking

*M. oryzae* conidia germinate predominantly from the apical cell, shortly after adhesion to the hydrophobic leaf surface by means of spore tip mucilage released from the apical compartment (Hamer et al., 1988; Saunders et al., 2010). Given that the apical cell is the sole reservoir for organelle inheritance by the appressorium, we decided to test whether this was a feature of that specific cell, or whether germination is the critical signal to enable organelle trafficking. We therefore carried out an exhaustive search for very rare conidia germinating from the blunt distal end in a fungal strain expressing Scad2-RFP and β-tubulin-GFP (Fig. 10). We observed that these conidia were still competent to make appressoria but that organelle trafficking occurred exclusively from the blunt germinating cell. Furthermore, in these cells, the apical cell underwent autophagy first (4h:30 min) followed by the middle cell (7h:30min). This suggests that the conidium has developmental plasticity to allow either terminal cell to germinate and become competent to undergo organelle trafficking, which then re-programmes the remaining cells for sequential autophagy and regulated cell death.

**Fig. 10.**
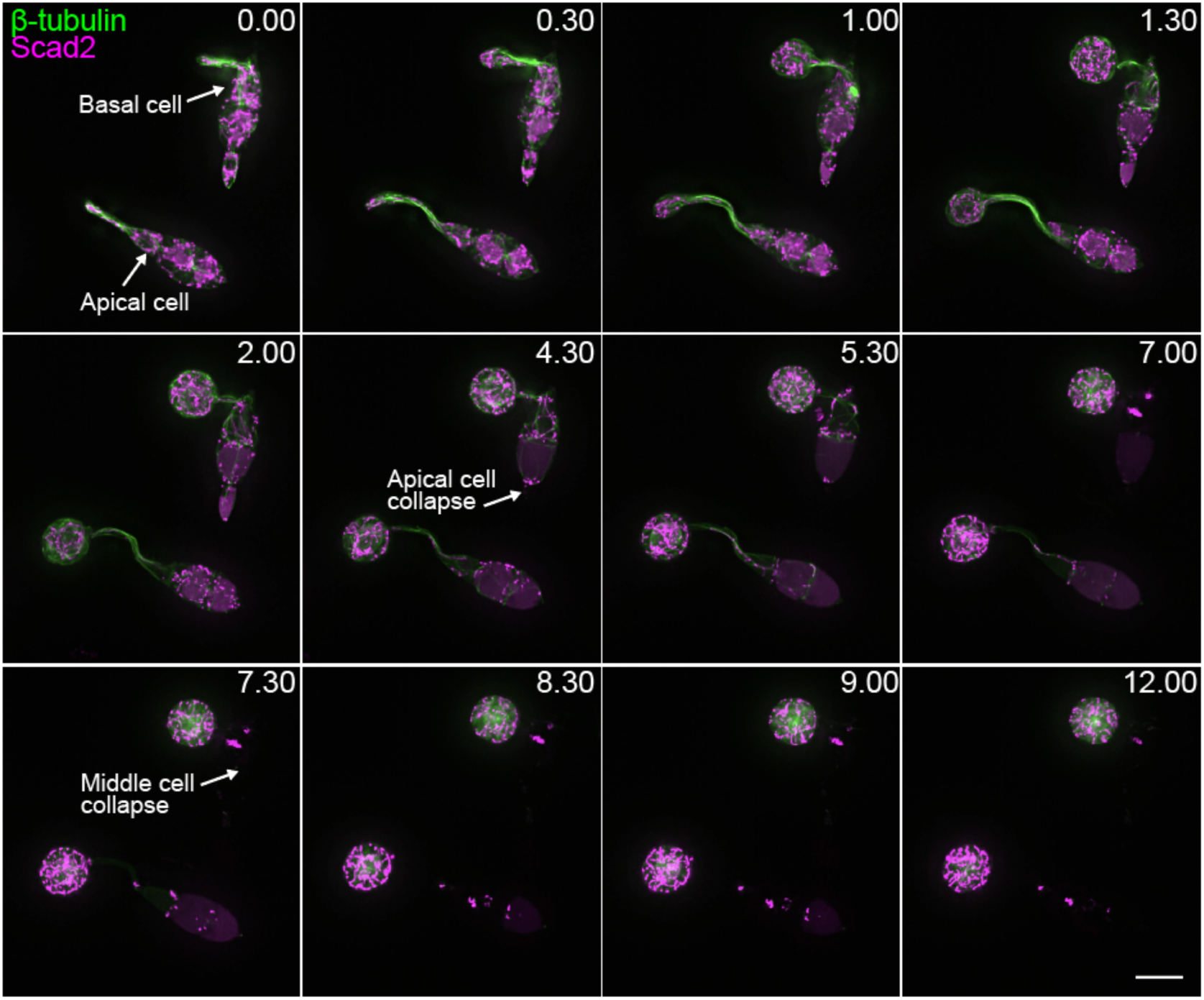
Germination site dictates cell fate and organelle inheritance in *M. oryzae* conidia. Time-series contrasting mitochondrial (Scad2-RFP) and microtubule (β-tubulin-GFP) dynamics and cell fate in *M. oryzae* conidia germinating from the apical versus basal cell. Micrographs represent maximum intensity projections of Z-series acquired at 0.25 μm intervals through the entire depth of the spores. Scale bar = 10 μm.

## Discussion

Appressorium development in *Magnaporthe oryzae* follows a tightly orchestrated morphogenetic process, governed by cell cycle checkpoints and requiring autophagy-dependent programmed cell death (Ryder et al, 2022; Eseola et al., 2021). In this study, we addressed a key question regarding plant infection by a pathogenic fungus. How is active fungal growth regulated during appressorium-mediated plant infection at the same time as programmed cell death? Given that many pathogenic fungi are faced with this developmental challenge, the underlying processes are likely to be of general importance to regulation of plant infection by fungi. In *M. oryzae* the unicellular appressorium penetrates the plant cuticle with a rigid penetration peg and all subsequent fungal growth derives from this single fungal cell, because the remaining pre-penetration stage fungal cells all undergo programmed cell death. This means that the appressorium must contain organelles and cytoplasm (originating from the conidium) from which subsequent organelle biogenesis and active fungal growth proceeds. To understand these processes, we set out to investigate the spatio-temporal control of organelle trafficking and autophagy during appressorium development using high-resolution 4D live-cell imaging of *M. oryzae* expressing constitutive and conditionally fluorescent tagged organelles.

The first major conclusion of this study is that organelles are actively trafficked from the conidium to the appressorium during its development. Trafficking occurs from the onset of conidial germination and is completed by 6h. We showed that normal organelle trafficking requires both actin filaments and microtubules, which mirrors observations in yeast and other fungi and likely involves the actin-associated motor protein myosin Myo2 (Li et al., 2018; Li et al., 2021; Pantazopoulou et al., 2014), in addition to long-range movement via microtubules (Dulal et al., 2021; Penalva et al., 2017; Steinberg and Schliwa, 1993). It has been reported that microtubule plus ends are directed towards the appressorium in *M. oryzae* (Dulal et al., 2021), consistent with a kinesin being necessary for organelle trafficking as reported in other filamentous fungi (Lin et al., 2016; Penalva et al., 2017). The second major conclusion of the study is that the germinated three-celled conidium is an open system that facilitates intercellular trafficking. This conclusion is evidenced by the observation that conditional activation of cytoplasmic paGFP results in its movement between individual conidium cells and into the germ tube and incipient appressorium. This shows that cytoplasm can freely diffuse between cells, consistent with septal pores being open during the onset of appressorium development. This occurs up to the point of appressorium cytokinesis, suggesting that organelles must also have the capacity to move between cells to the incipient appressorium. Organelle trafficking between fungal cells in hyphae is well known (Steinberg and Schliwa, 1993; (Bielska et al., 2014; Callejas-Negrete et al., 2025; Lin et al., 2016; Schuster et al., 2025) and can facilitate very long-range movement of organelles in fungal hyphae (Bowman, 2025). Septal pores in fungi can be closed by specialised structures called Woronin bodies, which occurs during the cell cycle (Shen et al., 2014) or as result of cell injury (Mamun et al., 2023). During appressorium development, completion of mitosis and nuclear migration triggers formation of an actomyosin contractile ring at the neck of the incipient appressorium and an asymmetric cytokinesis (Saunders et al., 2010b), in which the septal pore is closed by a Woronin body (Soundararajan et al., 2014). Our live cell imaging results confirmed that closure of the appressorium septum occurs by 6 h after germination, effectively sealing the appressorium from the collapsed conidium and germ tube after that time.

The third major conclusion emerging from this study is that each cell of the conidium undergoes a distinct developmental program. Using paGFP to conditionally monitor mitochondria within individual cells of the conidium revealed that the distal and middle cells of the conidium are programmed to undergo autophagy sequentially, starting with the distal cell. By contrast, all organelles in the appressorium originate from the apical conidial cell. If autophagy is inhibited, organelles remain intact in the distal and middle cells but still traffic from the apical cell to the appressorium. This means that although the conidium is an open system capable of trafficking contents from any of its cells during appressorium development, organelles are held in place within two of these cells. Moreover, although we focused on mitochondria as an example of an organelle requiring template-based biogenesis and therefore required to move between fungal cells (Bowman, 2025), our observations were mirrored in all of the organelle types we examined. Distinct spatio-temporal regulation of autophagy must therefore occur in distal, middle and apical cells of *M. oryzae* conidia, while only one of these cells is competent to traffic organelles. We reasoned that one likely signal to achieve three distinct developmental fates in each cell would be via cell cycle regulation. The nucleus of the apical germinating cell, for example, must pass into S-phase for appressorium development to be initiated (Osés-Ruiz et al., 2017; Saunders et al., 2010b; Veneault-Fourrey et al., 2006a). Similarly, nuclear division and daughter nucleus migration to the incipient appressorium must occur, via a G2-M phase checkpoint for the infection cell to mature and generate turgor (Veneault-Fourrey et al., 2006a), while mitotic exit is necessary for its competence to cause infection (Saunders et al., 2010a), which requires a further pressure-dependent S-phase checkpoint in the appressorium to trigger septin-dependent re-polarisation (Osés-Ruiz et al., 2017). Surprisingly, none of these developmental checkpoints are required to regulate organelle trafficking. The spatial control of organelle trafficking is therefore cell cycle-independent. Interestingly, this is consistent with recent evidence in yeast which also suggests that organelle inheritance during budding is cell cycle-independent (Li et al., 2021). What then is the signal to enable organelle trafficking to the appressorium? It seems likely, based on our observations, that spore germination signals the distinct developmental fate of each conidial cell with apical cells normally acting as the organelle reservoir. Rare conidia that germinate from the blunt distal end, strikingly, underwent the opposite pattern of inheritance, undergoing autophagy and regulated cell death first in the apical cell, followed by the middle cell, while it was the blunt distal cell that trafficked organelles to the developing appressorium, prior to its final collapse. This suggests a level of plasticity in conidial germination that still facilitates appressorium development, but which consequently necessitates an alternative sequence of autophagy-dependent cell death. As regulated cell death has recently been reported to require ferroptosis (Shen et al., 2020; Wengler and Talbot, 2025), this also suggests distinct spatial control of the final cell death signal must operate in each conidial cell.

Finally, a consequence of organelle movement to the incipient appressorium is that rapid *de novo* organelle biogenesis can occur prior to plant infection, thereby providing the infection cell with a reservoir for the extensive invasive hyphal growth that it must undertake upon entry into the host. Using a photoconvertible fluorescent reporter enabled the precise timing of organelle biogenesis to be determined, shortly after trafficking occurred and following septation of the appressorium (8h after germination). Volumetric analysis and conditional photoconvertible fluorescence microscopy also revealed that organelle biogenesis proceeds independently of conidial autophagy. This means that breakdown of organelles in the distal and middle cells of the conidium is not necessary for subsequent organelle biogenesis. This highlights again the distinct developmental fate of each conidial cell and the cell autonomous nature of organelle biogenesis in the appressorium. However, these observations raise further questions. What is the precise role of autophagic recycling from the conidium, for example, if it is not strictly necessary to fuel a process as intrinsic to growth as organelle biogenesis? Given that autophagy in *M. oryzae* is essential for plant infection (Veneault-Fourrey et al., 2006), the obvious answer is that autophagy must be required for appressorium turgor generation. Turgor is produced by an influx of water into the appressorium as a result of glycerol generation in the cell (de Jong et al., 1997). The presence of a melanin layer in the appressorium cell wall prevents efflux of glycerol and other solutes, allowing the appressorium to generate turgor (de Jong et al., 1997). Incipient cytorrhysis assays of Δ*atg8* and Δ*atg1* mutants show, for example, that they generate lower turgor compared to the isogenic wild-type strain Guy11 (Kershaw and Talbot, 2009). So, autophagic recycling may be necessary for glycerol synthesis to high concentrations, although this has not been directly confirmed biochemically. It also seems likely, however, that autophagy acts as a developmental checkpoint for appressorium morphogenesis, especially given the precise temporal order of regulated cell death revealed in this study. Consistent with the developmental checkpoint hypothesis, repolarisation of the appressorium does not occur in autophagy mutants, although their precise role of autophagy in the regulation of septin assembly at the appressorium pore– a pre-requisite for re-polarisation – has not been investigated.

In summary, our study has revealed that synchronous spatio-temporal control of autophagy and organelle trafficking occurs during appressorium morphogenesis by the rice blast fungus. The pathogen has therefore evolved the ability to simultaneously activate turgor-dependent appressorium-mediated plant infection, resulting in very rapid fungal growth, while undergoing programmed cell death and autophagic recycling. This remarkable ability requires completely distinct developmental programmes to operate simultaneously in connected fungal cells in order to facilitate plant infection.

---

**Supplementary Fig. S1.**
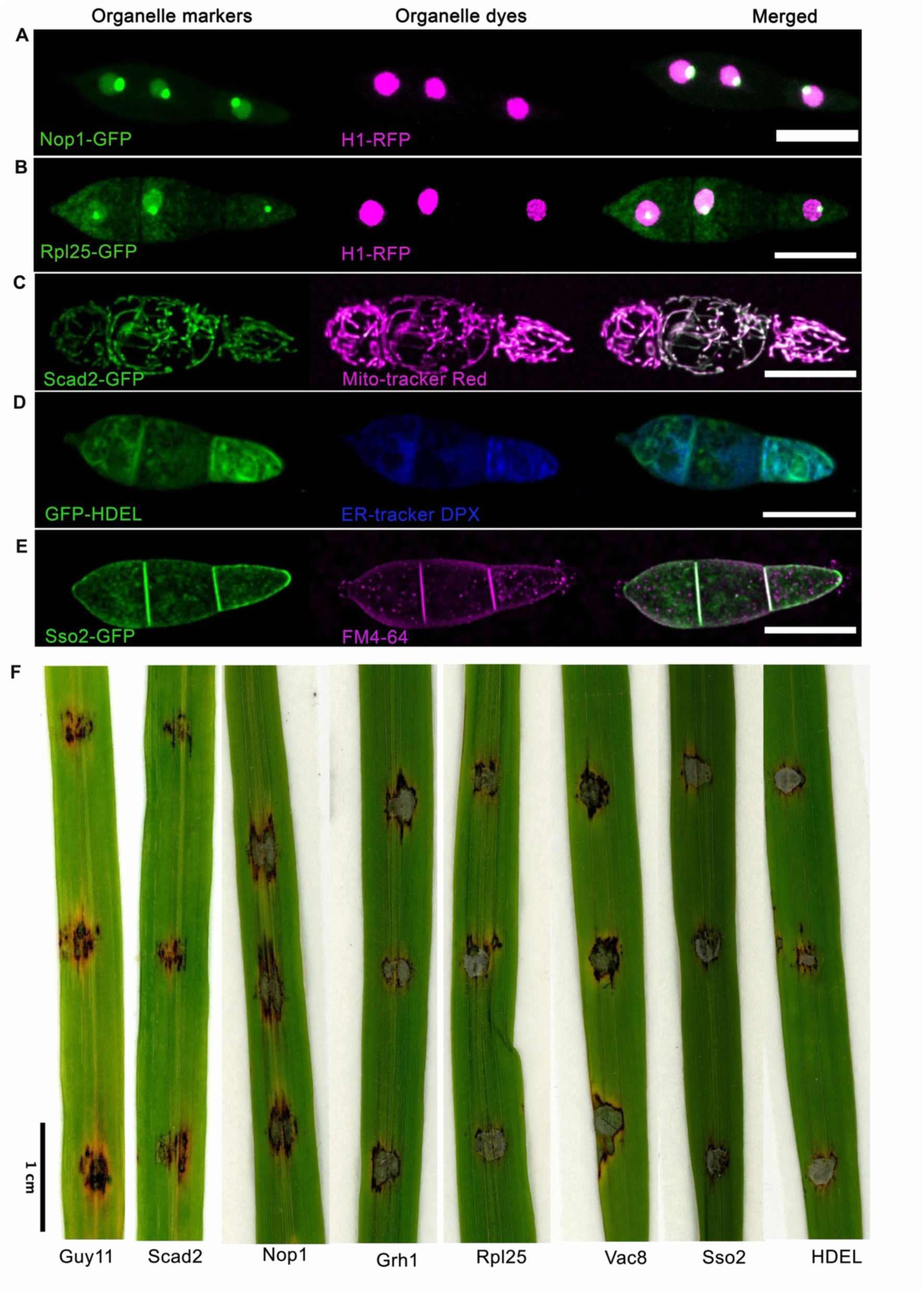
Co-localisation of fluorescent fusion proteins with organelle-specific markers and confirmation of the ability of all generated strains to cause rice blast disease. Micrographs of (**A**) *M. oryzae* conidium co-expressing nucleolar Nop1-GFP and nuclear H1:RFP. (**B**) Conidium co-expressing nuclear histone marker; H1-RFP and 60S ribosomal marker; Rpl25-GFP. Accumulation of the green signal as a punctate object colocalised to a subregion of the red fluorescence of the nucleus histone; H1-RFP. (C) Conidium of *M. oryzae* expressing mitochondrial marker; Scad2-GFP co-stained with MitoTracker Red CMXRos dye. The dye perfectly colocalised with the elongated tubules of Scad2-GFP confirming that Scad2-GFP is localised to the mitochondria. (D) *M. oryzae* conidium expressing plasma membrane marker; Sso2-GFP co-stained with the liphophilic dye FM4-64. The green fluorescence signal of Sso2-GFP perfectly co-localise with the red fluorescence of the FM4-64 dye to label the periphery and septa of the conidium. (E) Conidium of *M. oryzae* expressing the ER marker; GFP-HDEL counterstained with ER specific stain; ER-Tracker DPX. The merged image confirmed that GFP-HDEL is localised in the ER. The micrographs represent the maximum intensity projection of a z-stack series. All red images were false-colour adjusted to magenta using ImageJ (US National Institute of Health). The scale bars represent 10 µm. (F) Images of rice leaves showing infection lesions formation four days post-inoculation with conidia of *M. oryzae* strains expressing single copy integration of individual organelle markers used in this study. The presence of lesions on the rice leaves indicates that the GFP-tagged organelles are fully pathogenic.

**Figure S2.**
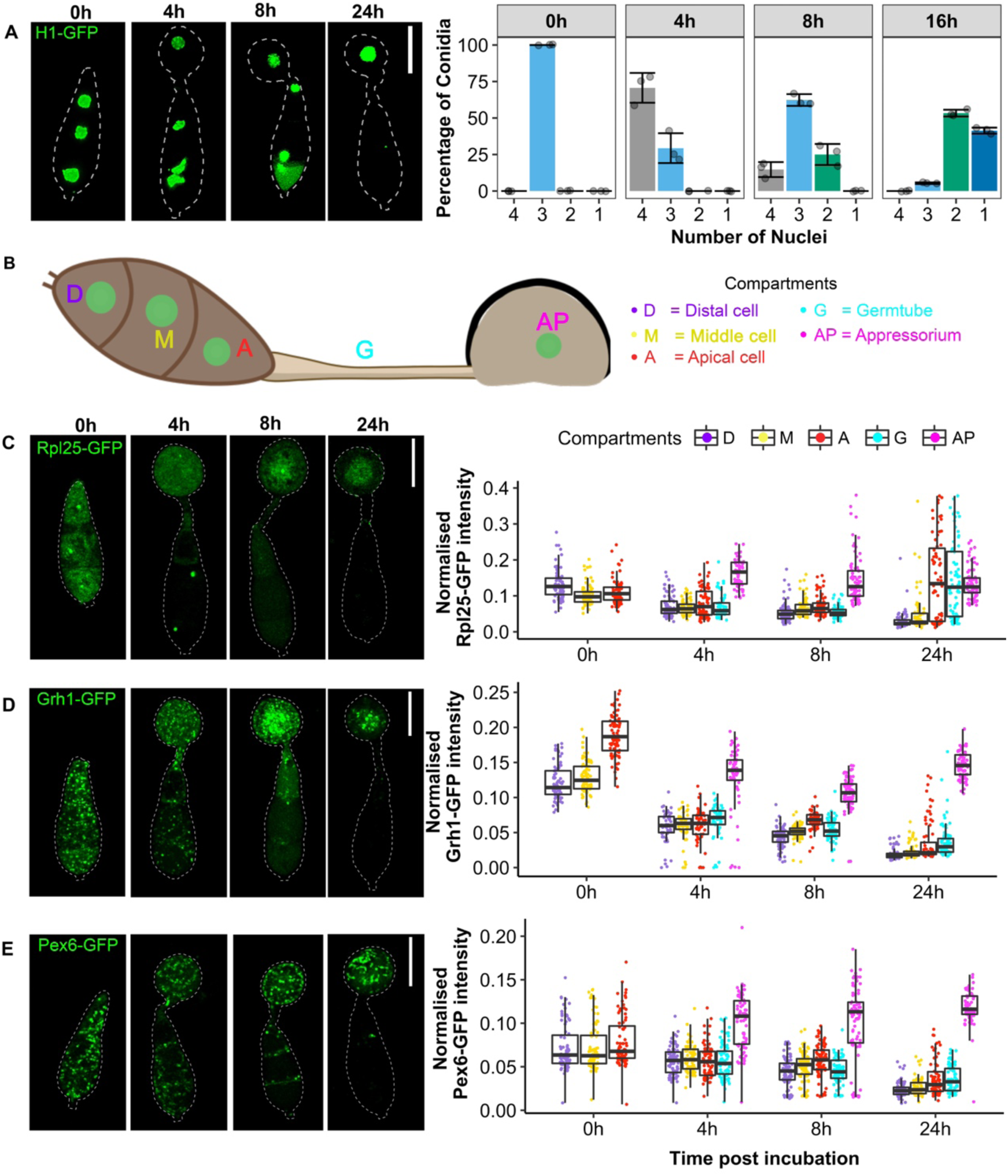
Sub-population of organelles are trafficked to the appressorium during development. (**A**) Representative fluorescence micrographs showing nuclear dynamics and a corresponding bar chart showing nuclei dynamics during appressorium development of *M. oryzae.* (B) Image illustrating the labelling of the different cellular compartments of *M. oryzae*, which was used to quantify fluorescence intensity. (C) Micrographs of *M. oryzae* conidia expressing ribosomal marker; Rpl25-GFP and the corresponding boxplots to show ribosomes dynamics (Supplementary Video S6) in the conidia of *M. oryzae* at specific time of appressorium development. (D) Micrographs of *M. oryzae* conidia expressing early Golgi marker; Grh1-GFP and the corresponding boxplots to show early Golgi dynamics (Supplementary Video S7) at indicated time points of appressorium development. (E) Micrographs of *M. oryzae* conidia expressing peroxisome marker; Pex6-GFP and the corresponding boxplots to show peroxisome dynamics (Supplementary Video S8) at indicated time points of appressorium development. The images are maximum intensity projections of Z-stacks taken at 0h, 4h, 8h, and 24h after incubation. The white dashed line outlines the cell boundary drawn with ImageJ; the scale bar represents 10 µm. Boxplots shows organelle population within each cell compartments estimated by the mean normalised quantified fluorescence intensity of each organelle marker. The different coloured points indicate each conidium compartment (N = 25 conidia, 3 replicates). The horizontal line of the boxplot is the mean of three replicates and the box represents the quantile, the error bar represents the standard deviation. (E) Micrographs of *M. oryzae* conidia expressing peroxisome marker; Pex6-GFP and the corresponding boxplots to show peroxisome dynamics (Supplementary Video S8) at indicated time points of appressorium development. The images are maximum intensity projections of Z-stacks taken at 0h, 4h, 8h, and 24h after incubation. The white dashed line outlines the cell boundary drawn with ImageJ; the scale bar represents 10 µm. Boxplots show organelle population within each cell compartments estimated by the mean normalised quantified fluorescence intensity of each organelle marker. The different coloured points indicate each conidium compartment (N = 25 conidia, 3 replicates). The horizontal line of the boxplot is the mean of three replicates and the box represents the quantile, the error bar represents the standard deviation.

**Figure S3.**
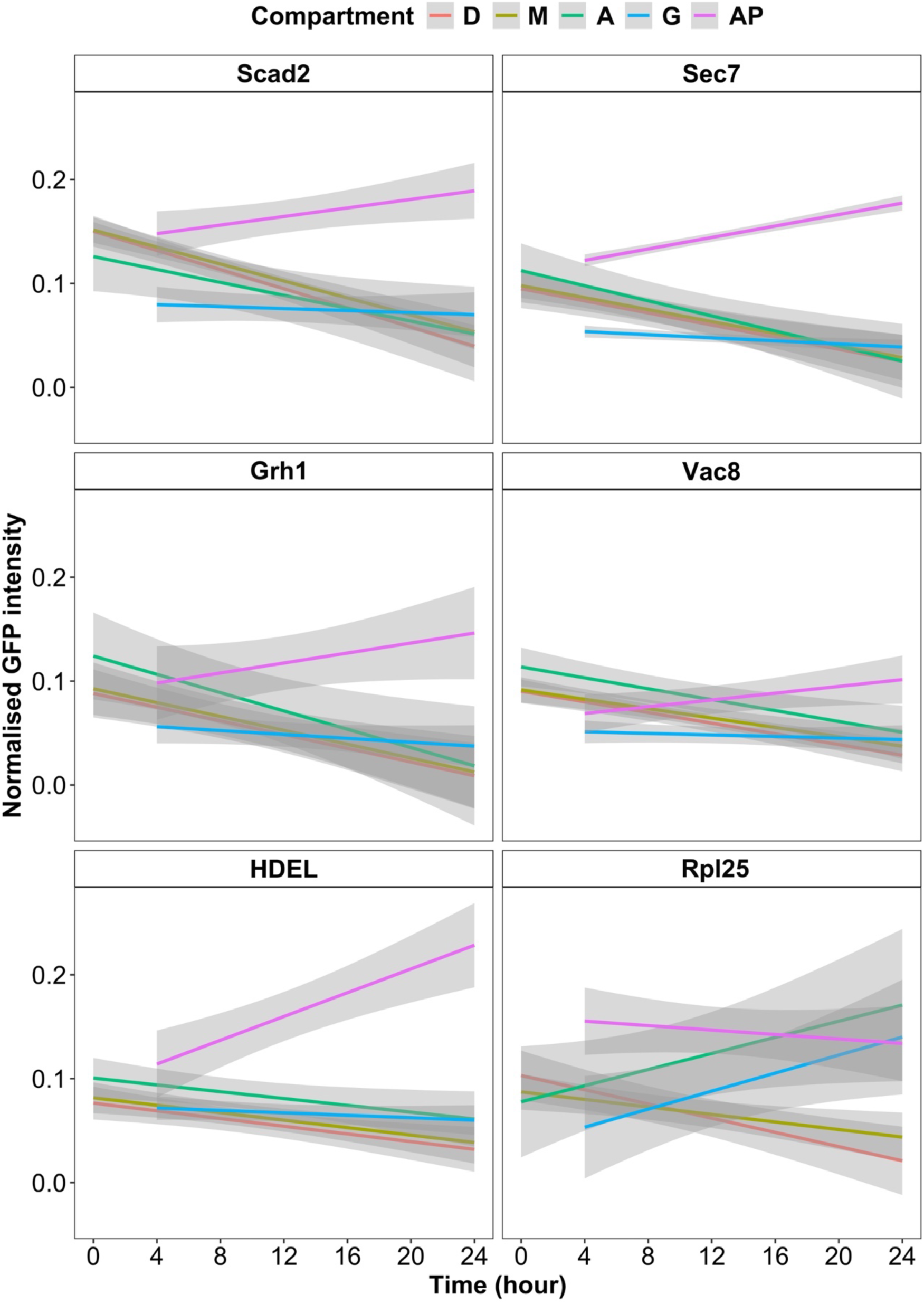
Simple linear regressions of fluorescence intensity on time in given cell regions during infection-related-development. The shaded area is the standard error. The line graph was calculated from the means of three replicates (N = 25 conidia) using a general linear model in R.

**Figure S4.**
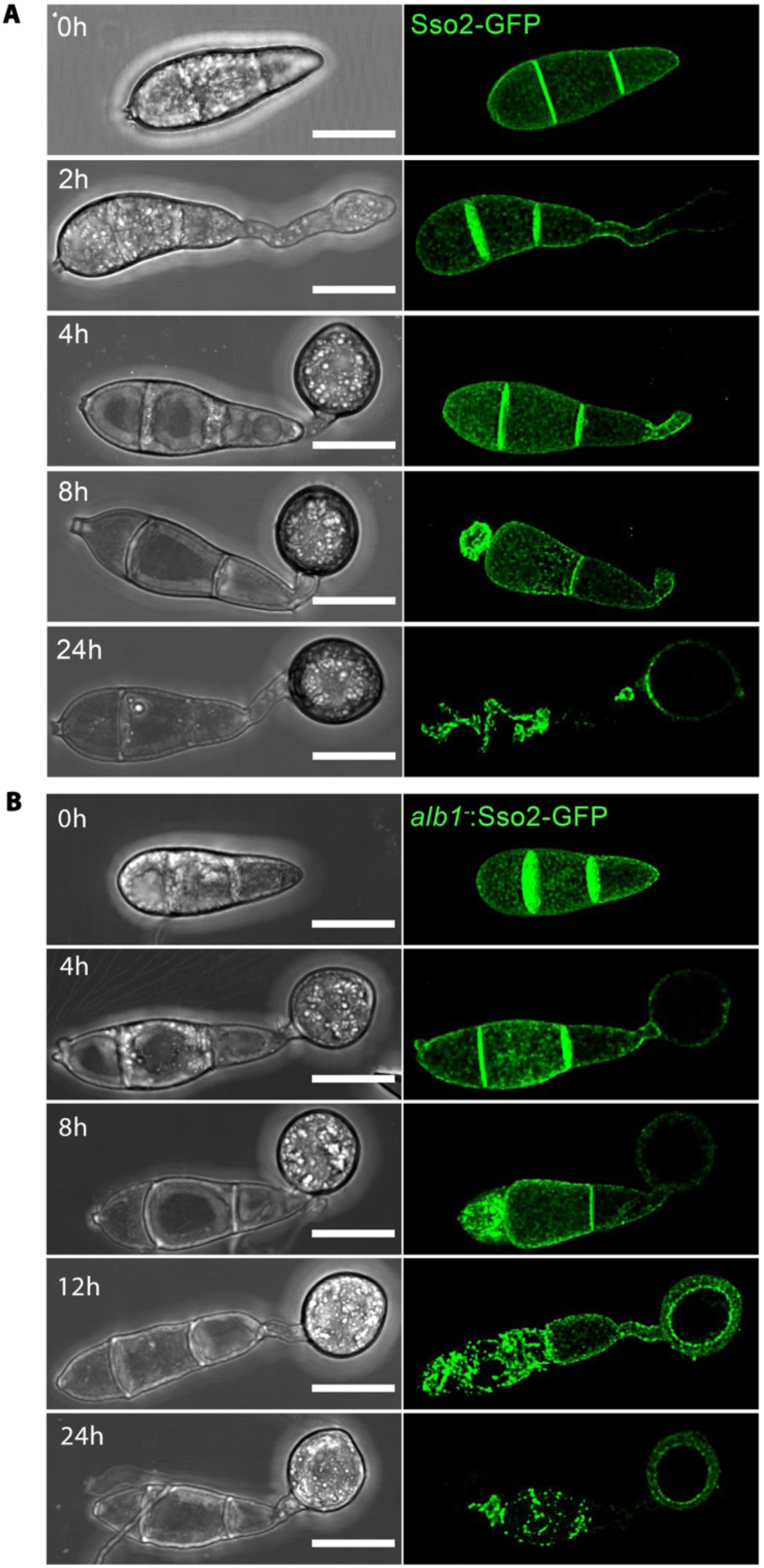
Organisation of the plasma membrane dynamics during appressorium development of *M. oryzae*. (A) Representative micrographs of *M. oryzae* conidia of Guy11 strain expressing plasma membrane marker Sso2-GFP. The marker labels the conidia cell periphery and septa. From 8h, the plasma membrane of conidia disintegrated from distal, middle and apical cells. At 24h, Sso2-GFP faintly labels the appressorium cortex. (B) Representative micrographs of *M. oryzae* conidia of albino mutant *alb1-* expressing Sso2-GFP. The marker localised at the appressorium cortex and showed integrity of the conidial plasma membrane during early appressorium development. The conidial plasma membrane broke down between 8 and 24 h starting with the distal cell, then the middle and apical conidial cells. At 24h, the expression of Sso2-GFP was clearly visible at the appressorium cortex. Micrographs shown are maximum intensity projections of the *Z*-stacks series; the scale bar represents 10 µm

**Figure S5.**
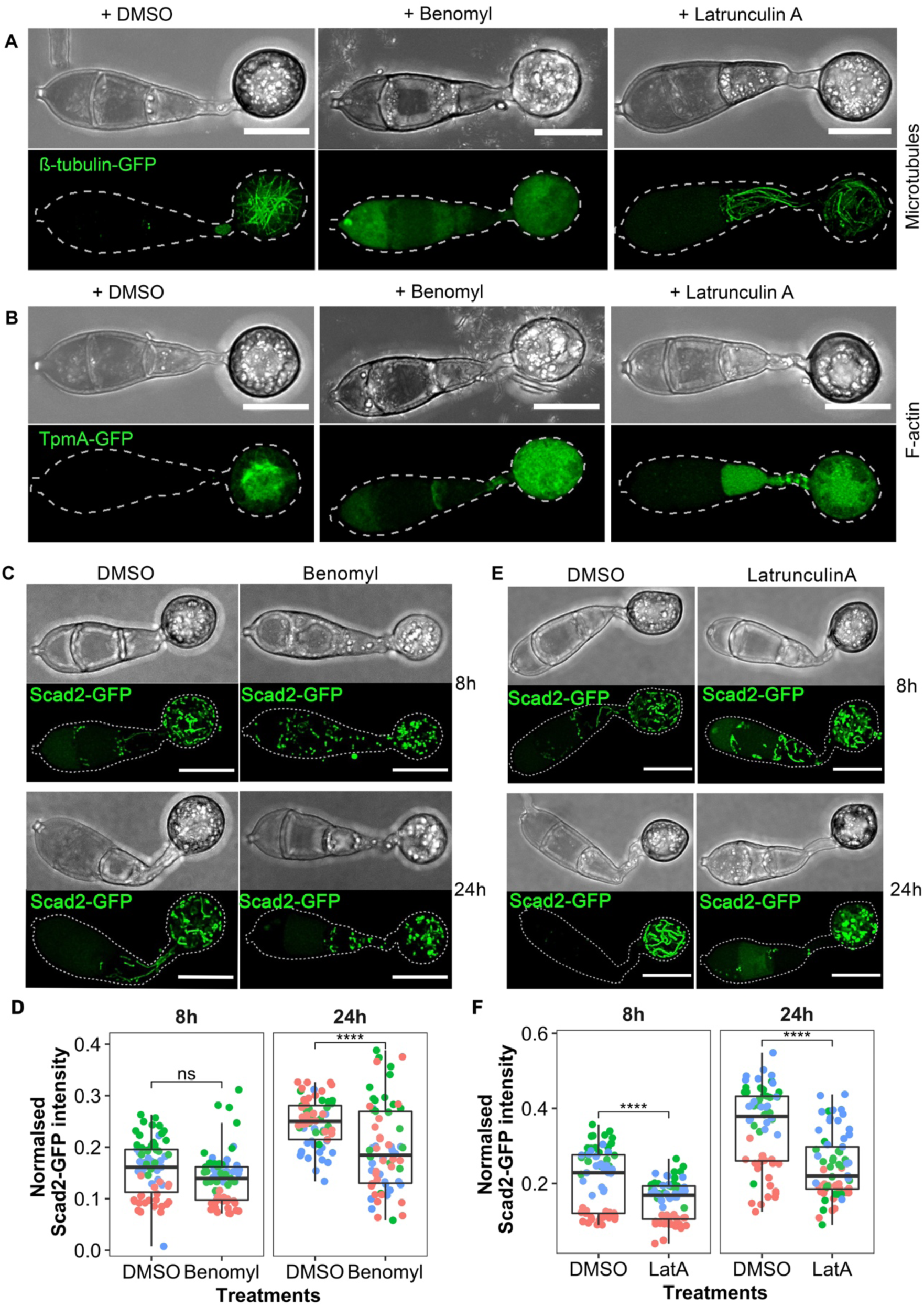
Intact microtubules and actin filaments are required for mitochondria transport in the appressorium of *M. oryzae*. (A) Micrographs to show the effect of benomyl and latrunculin-A on microtubules visualised using ß-tubulin-GFP. Benomyl treatment leads to complete depolymerisation of microtubules, while latrunculin-A treatment did not disrupt microtubules. (B) Micrographs show the effect of benomyl and latrunculin-A treatments on F-actin organisation visualised using a tropomyosin marker, TpmA-GFP. Both benomyl and latrunculin-A treatments lead to mislocalisation of F-actin organisation at the centre of appressorium. DMSO treatment did not affect microtubules or F-actin organisation. In addition, benomyl treatment prevented melanisation of the appressorium cell wall. (C) Micrograph showing the fragmentation and uneven distribution of mitochondria; Scad2-GFP in germinating conidia treated with 10 µg mL^-1^ of the microtubule inhibitor benomyl and DMSO control. (D) Boxplot showing Scad2-GFP intensity in the appressorium of benomyl and DMSO treated samples. The treatments were added at 4 hpi and images acquired at 8hpi and 24 hpi. The boxplot was calculated from 25 conidia per experiment with three replications of the experiment. (E) Micrograph showing depolymerisation of actin filaments using 10 µM latrunculin-A and its effect on mitochondria as visualised by marker Scad2-GFP. Scale bar represents 10 µm. (F) Boxplot showing Scad2-GFP intensity in the appressorium of latrunculin-A and DMSO treated samples. The treatments were added at 4 hpi and images acquired at 8hpi and 24 hpi. Boxplots were calculated from 25 conidia per experiment with three replications and significance difference calculated using unpaired t-test (**** = P<0.0001).

**Figure S6.**
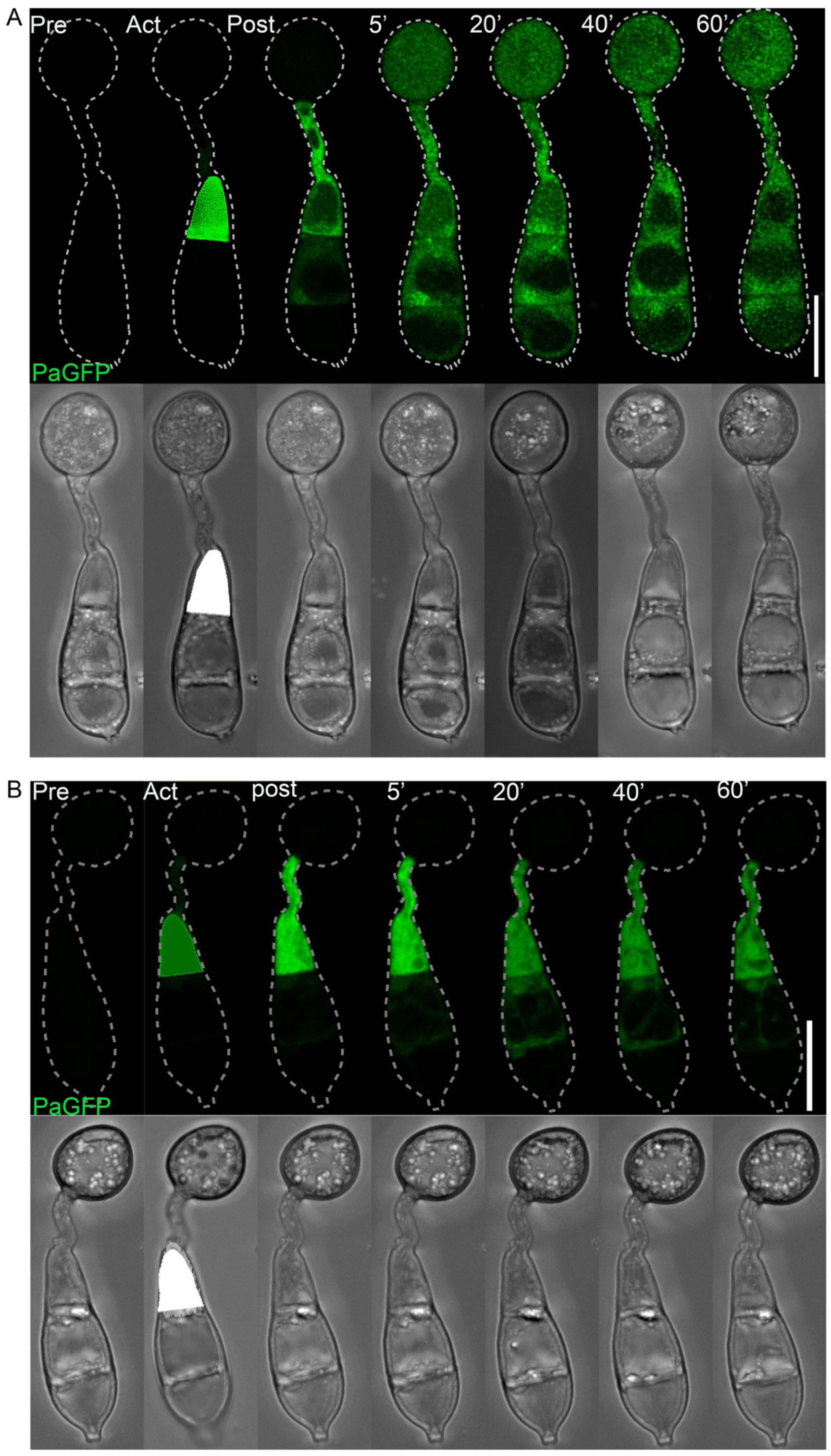
Photoactivation experiment reveals the timing of cytokinesis at the appressorium neck. Conidia expressing paGFP were incubated on a hydrophobic coverslip before photoactivation. Before photoactivation, paGFP expression was not visible. (A) Micrograph of confocal images of conidia photoactivated after 4h post-incubation. After a short pulse with the 405-nm laser, fluorescence appeared to traffic into adjacent cells. The displayed stilled frames are maximum intensity projections of the *Z*-stack series taken from (Supplementary Video S15). (B) Micrograph of confocal images of conidia photoactivated at 6h after germination. After photoactivation, fluorescence appeared to be restricted to the appressorium collar. The displayed stilled frames are maximum intensity projections of the *Z*-stack series taken from (Supplementary Video S16). The dotted line denotes the cell outline and was performed in ImageJ (US National Institutes of Health). Time is displayed in minutes, and the scale bar represents 10 µm.

**Figure S7.**
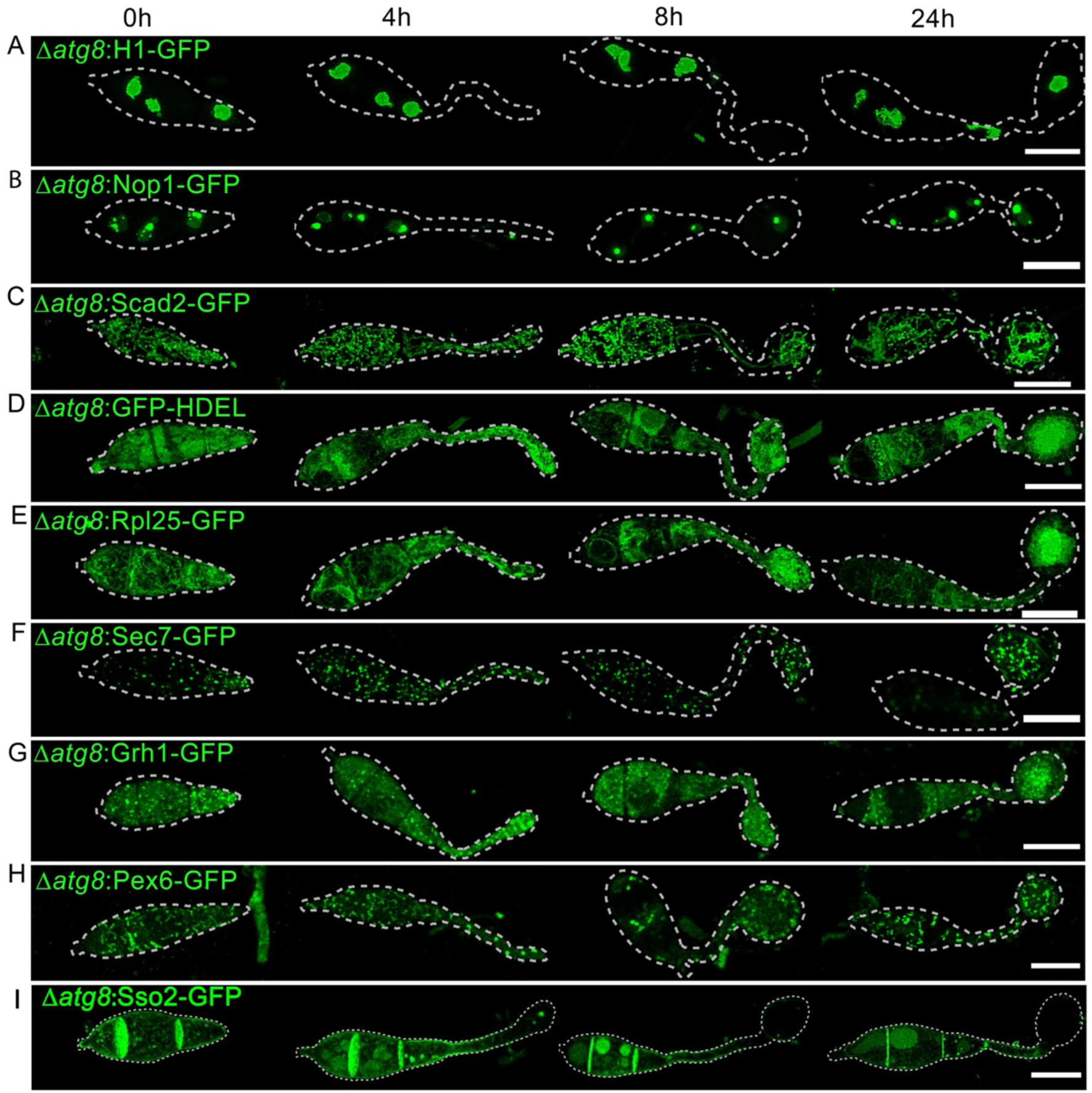
Organelle degradation is dependent on autophagy. Conidia were harvested from *τιatg8* mutant expressing (A) nuclei marker; H1-GFP (B) nucleoli marker; Nop1-GFP, (C) mitochondria marker; Scad2-GFP (D) endoplasmic reticulum marker; GFP-HDEL, (E) ribosome marker; Rpl25-GFP, (F) trans-Golgi marker; Sec7-GFP, (G) early Golgi marker; Grh1-GFP, (H) peroxisomes marker; Pex6-GFP, and (I) plasma membrane t-SNARE marker, Sso1-GFP. Conidia were incubated on hydrophobic coverslip and observed at 0, 4, 8, and 24 hpi by laser confocal microscopy. Micrographs are maximum projection of *Z-*stack series captured at the specified times during appressorium development. Scale bar represents 10 µm.

**Figure S8.**
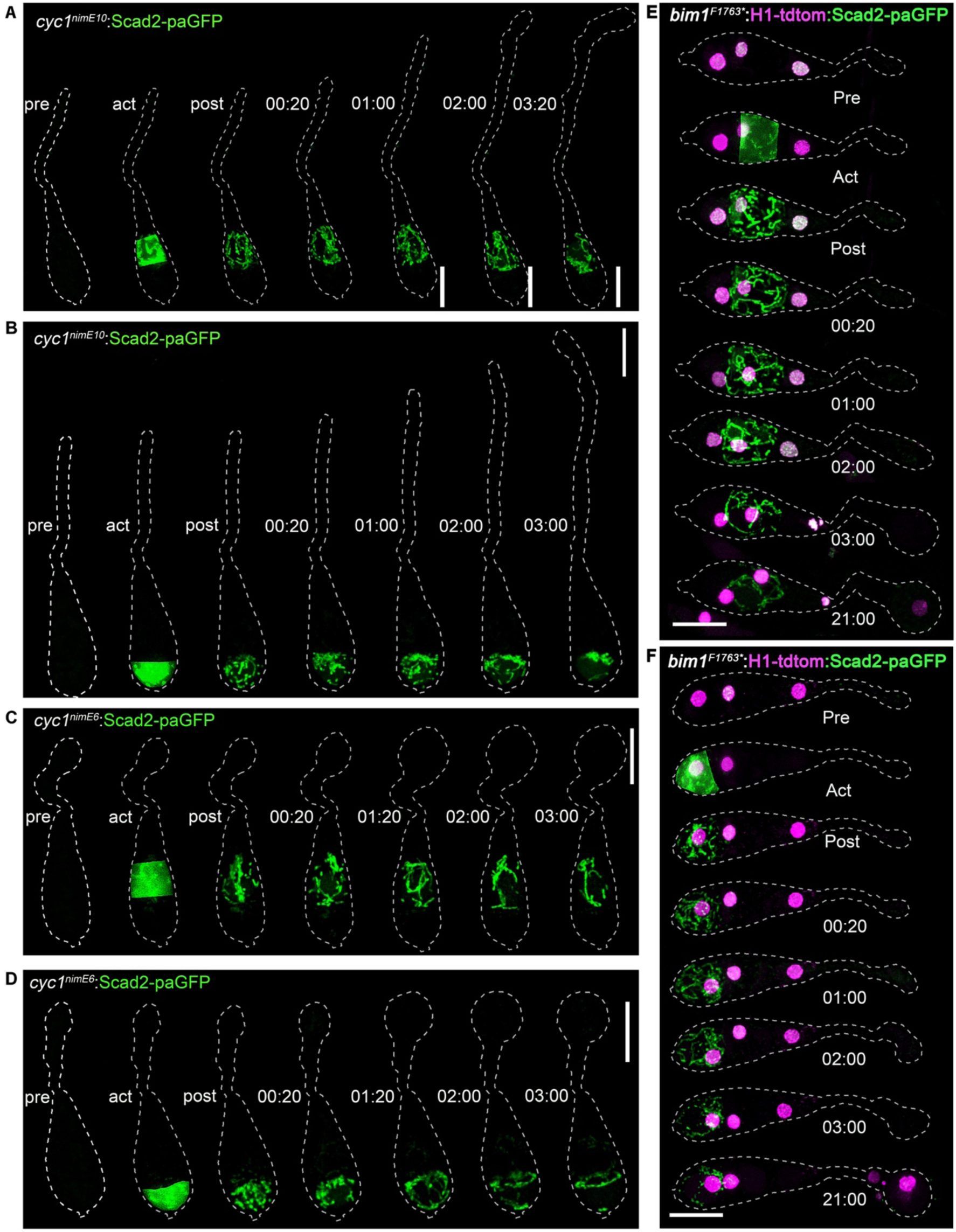
Organelle trafficking is not regulated by cell cycle checkpoints in M. oryzae. Micrographs of conidia of conditional S-phase mutant expressing *cyc1^nimE10^*:Scad2-paGFP. After photoactivation of the (A) middle and (B) distal conidium cell, fluorescent mitochondria were restricted to the individual cells and no movement was observed into the adjacent cells (C) Micrographs of conidia of conditional G2-phase mutant expressing *cyc1^nimE6^*:Scad2-paGFP, after photoactivation of the middle and (D) distal conidium cell, fluorescent mitochondria showed no movement to the adjacent cells (E) Micrograph of conidium of conditional mitotic mutant photoactivated in the middle and (F) distal cells at 3h post-incubation. After photoactivation, fluorescent mitochondria in the showed no movement to the adjacent cells and the conidia contents were not degraded. The conidia of the coniditional mutants were incubated on hydrophobic coverslips at 30°C before photoactivation and live-cell imaging in a preheated microscope chamber. The dotted line outlines the cell and was performed in ImageJ. Time is displayed in hours:minutes post activation, and the scale bar represents 10 µm.

**Figure S9.**
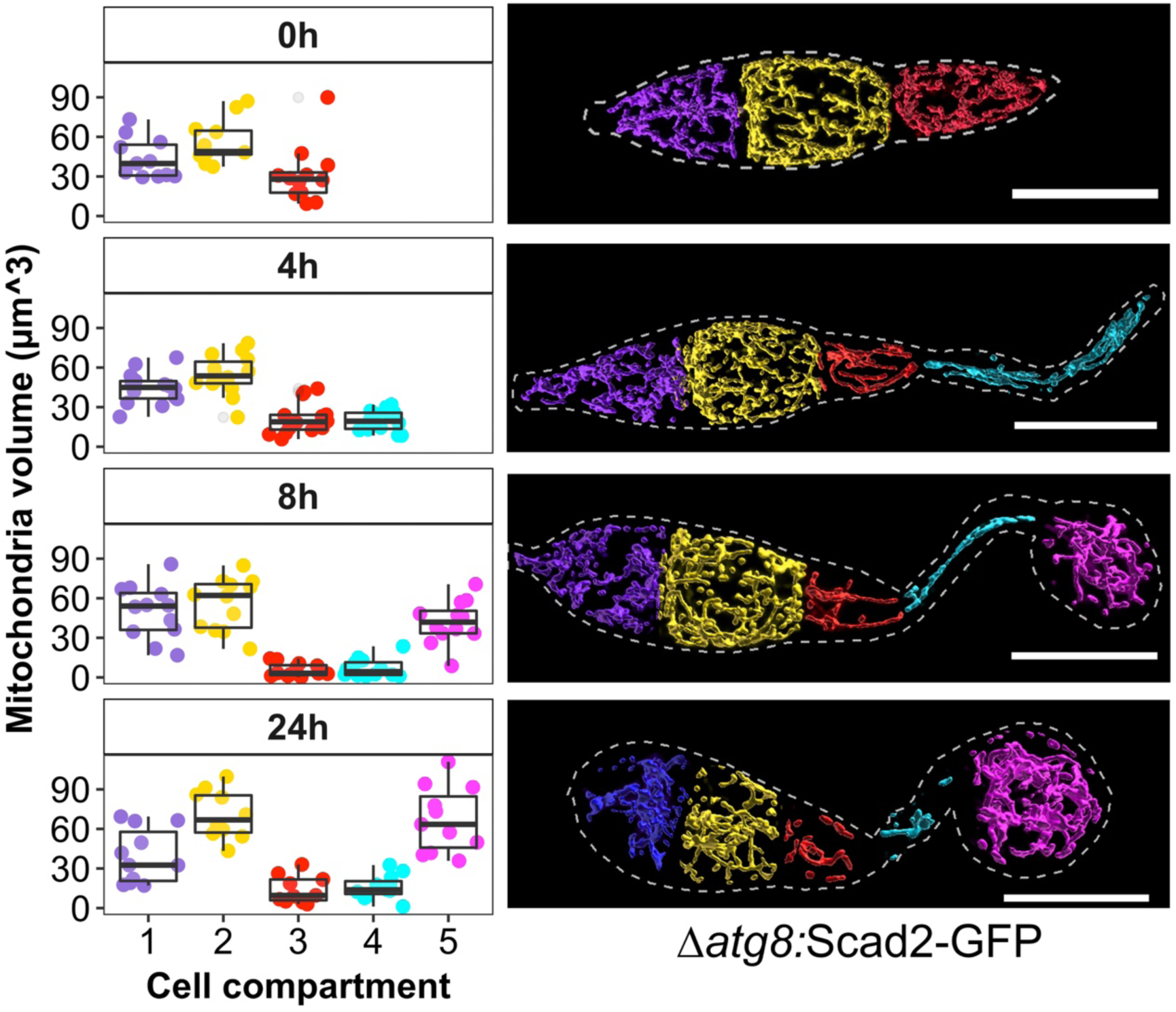
Volumetric analysis of intact mitochondrial during appressorium development of *M. oryzae*. Boxplot and corresponding micrographs showing the volume of intact mitochondria within conidia of a Δ*atg8* null mutant of *M. oryzae* during appressorium development at 0, 4, 8, and 24 h. The different colours in the plot represent each conidium cell (N= 12 conidia). The scale bar represents 10 µm. Image rendering was performed in Imaris.

## Materials and methods

### Fungal strains, growth condition and appressorium assay

All fungal strains generated in this study were generated in the wildtype rice pathogenic strain Guy11 (Leung et al., 1988) and an isogenic autophagy-deficient *Δatg8* mutant (Kershaw and Talbot, 2009). The cultures were routinely grown on complete medium (CM) agar (Talbot et al., 1993) at 25°C under 12-hour light and dark cycle for nine days. Conidia from nine-day old cultures grown on CM plate were harvested by flooding the plate with sterile de-ionised water and using a sterile spatula to brush the culture surface to release the conidia gently. The conidial suspension was filtered through sterile Miracloth (Calbiochem) and centrifuged at 10 000 g for 5 min at room temperature. A 50 µL conidia suspension was placed onto hydrophobic glass coverslip at a concentration of 5 x 10^4^/mL to monitor conidial germination and appressoria development over 24 h duration at an interval of 4 h. All the strains generated in this study are stored as desiccated filter stocks and are available from the laboratory of NJT at The Sainsbury Laboratory, Norwich.

### Identification of *M. oryzae* organelle’s homologues

The homologue of each organelle described in this study was obtained by screening published organelle-specific markers described for *Saccharomyces cerevisiae, Neurospora crassa, Zymoseptoria tritici, Aspergillus nidulans* and was used in a blast search for the putative homologues in *M. oryzae*. These sequences were used to identify *M. oryzae* homologs using the BLAST function in EnsemblFungi (http://fungi.ensembl.org/Multi/Tools/Blast). The gene identification number for each organelle is provided in Supplementary Table S1.

### Construction of GFP gene fusion plasmids, fungal transformation, and determination of integrated GFP copy numbers

A 1.5 kb fragment upstream of individual organelle gene amplified together with the gene coding region. The PCR was performed using primers listed in Supplementary Table S2. The amplified DNA fragment together with a 1.5 kb fragment of enhanced green fluorescent protein GFP containing a TrpC terminator was cloned into pPscBAR vector using an infusion cloning kit from Takara Bio. To construct a vector containing A photo-activatable variant of GFP, a 750 bp PCR fragment of PAGFP was amplified from plasmid pPAGFP-C1 (Addgene plasmid # 11910). The PAGFP fragment was fused to the Trpc-T fragment in a second PCR reaction to obtain a single 1.5 kb amplicon. The 1.5kb fragment was cloned together with each specific organelle genomic DNA fragment using In-fusion (Takara) recombination cloning (Krappmann et al., 2006). The resulting plasmid DNA was transformed into protoplasts of the wild-type strain Guy11 or an autophagy-deficient mutant Δ*atg8*. All transformants were selected in the presence of 150 mgmL^-1^ glufosinate-ammonium (PESTANAL, Sigma-Aldrich). Genomic DNA was extracted from positive transformants expressing GFP. A 1ngµL^-1^ DNA concentration was sent to iDNA genetics Ltd (Norwich Research Park) to estimate the number of GFP insertions by quantitative real-time PCR according to (Bartlett et al., 2008). Transformants with a single GFP insertion were used in this study.

### Construction of photoactivatable mitochondria marker; Scad2-paGFP plasmid

A 3.1 kb region of the *SCAD2* encoding gene representing the coding sequence and 1.5 kb of promoter was amplified from plasmid pAE001 using primer pairs (Supplementary Table S2) with Phusion Taq polymerase. The PCR profile consisted of an initial denaturation step at 98°C for 30 s, followed by 35 cycles of denaturation at 98°C for 10 s, annealing at 67°C for 30 s, elongation at 72°C for 2 min and a final elongation step at 72°C for 7 min. The 3.1 kb amplicon was excised and gel purified. The purified fragment together with paGFP-trpC fragment were introduced into a linearized pscSUR cloning vector using the In-fusion (Takara) recombination cloning to give plasmid pAE010. Positive clones were selected by PCR and analysed by DNA sequencing.

### Appressorium development and rice infection assay

Conidia from nine days old culture grown on CM plate were harvested by flooding the plate with sterile de-ionised water and using a sterile spatula to gently brush the culture surface to release the conidia. The conidia suspension was filtered through a sterile Mira cloth (Calbiochem) and centrifuged at 10 000 *g* for five minutes at room temperature. For appressorium development, conidia were incubated on a hydrophobic glass coverslip (Menzel-Gläser) before visualisation. For plant infection assay, the conidia pellet was resuspended in 0.2% gelatine and used at a concentration of 1 x10^5^ conidia mL^-1^. A 20 µL aliquot of conidial suspension was inoculated onto detached leaves from a 3-week old susceptible rice cultivar CO-39 (Valent and Chumley, 1991) placed on 1% Agarose gel (Sigma). The samples were routinely incubated at 24°C with 12-hour light and dark cycle. Lesions were monitored and recorded five days post-inoculation.

### Treatment with inhibitory drugs

Benomyl and Latrunculin A were used at a concentration of 10µgml^-1^ and 10µM, respectively. The stock of the depolymerising microtubule drug, benomyl, was prepared by dissolving 10 mg benomyl (Methyl 1-(butylcarbamoyl)-2-benzimidazole carbamate) in 1 mL dimethylsulfoxide (DMSO). The stock of actin polymerisation inhibitor Latrunculin A (Sigma) was prepared by dissolving in DMSO. The inhibitors were added at 4 hpi by replacing the water of the incubated conidia with either treatment. The effect of the drug on organelle trafficking during appressorium development was examined at 4 and 20 h after treatment.

### Fluorescent cell markers

The conidia expressing GFP-tagged organelle marker were co-localised with organelle-specific fluorescent dyes. The mitochondria were stained with MitoTracker® Red CM-H2XRos (Life Technologies, Thermofisher cat. No. M7513) at a final concentration of 25 µM for 20 mins. The nucleus was stained with DAPI at a concentration of 1µgL^-1^. The nucleolar marker was co-localised with a strain expressing red fluorescent nuclear marker, H1:RFP, to visualise the nucleolus. Cell tracker blue was used at a concentration of 10 µM to visualise the vacuoles. The endoplasmic reticulum was stained for 10 mins with ER-Tracker Blue-White DPX (Thermonfisher, E12353) at a concentration of 20 µM. The plasma membrane probe, N-3-triethylammoniumpropyl-4-p-diethylaminophenylhexatrienyl-pyridinium-dibromide (FM4-64; Invitrogen, Karlsruhe, Germany), was used at a final concentration of 20 µM. BODIPY TR Ceramide (Thermofisher, D7540) was used to stain the Golgi bodies. Conidia were incubated with fluorescence dyes for 15 to 30 min before imaging. The excitation and emission of each fluorescent dye were performed according to manufacturer’s instructions.

### Laser confocal microscopy

All images were acquired using an inverted confocal laser microscope Leica TCS SP8X equipped with Leica DMi8 S module (Leica Microsystem Inc. Buffalo Grove, IL, USA). The laser confocal microscope is coupled with 63x/1.4 numerical aperture oil immersion objectives with a camera and Leica Application Suite X (LAS X) software. All time-lapse live-cell images were captured using an x2.3 optical zoom (1024 x 512 frame) at 5 min intervals, and single *Z-*stacks images were captured at 4x optical zoom (512 x 512 frame). Samples expressing GFP were excited using a 488 nm argon laser and acquired at the optimal detection emission of 500-530 nm. The pinhole was adjusted to 1.5 Airy discs for the argon laser at 488 nm to enable sampling with minimal laser percentage. For monitoring the dynamics of individual GFP-tagged organelles, conidia were incubated on hydrophobic coverslips with a 0.17mm thickness (Menzel Gläser #1.5) and imaged at 0, 4, 8, 12, and 24 h post-incubation. Raw microscope images were used for fluorescence quantification. Images for figure presentation were 3D deconvolved with the LIGHTNING adaptive function (now known as STELLARIS) on the SP8X Leica microscope (Leica Microsystem Inc. Buffalo Grove, IL, USA) to obtain super-resolution images.

### Fluorescence recovery after photobleaching FRAP and Photoactivation

The LASX Live Data Mode tool on the Leica SP8X was used to set up three confocal experiment jobs with the same setting parameter as the first window. The middle window was modified to define bleaching or photoactivation parameters, while the third window contains the same parameter as the first to collect the data after photobleaching with a 5 min interval between frames. A 405 nm laser was used at 70 % to bleach the defined ROI for 100 sec to monitor the recovery of the GFP signal after bleaching. The activation of PaGFP was carried out as described by (Patterson and Lippincott-Schwartz, 2002). The defined ROI was exposed to 70% 405 nm laser for 5 sec. The pre, post bleach and photoactivation data were acquired by excitation at 488 nm and emission at 500-530 nm. The acquired time-lapse image raw data were used for fluorescent intensity measurement while images for figure presentation were 3D deconvolved with the LIGHTNING adaptive function and display adjusted for brightness.

### Image data analysis

Confocal images captured as a Z-stack series were converted to a maximum projection. A shape-tool in ImageJ software (National Institute of Health) was used to draw an outline around each cell, and the region of interest was saved as a zip file. The region of the cell to be quantified was drawn with ImageJ, and fluorescence intensity was automatically quantified using the python script; FluorSeg==0.0.26.dev0 (Dan MacLean, 2021) installed to jupyter notebook. Normalised fluorescence intensity was calculated by dividing the fluorescence intensity by cell area. The relationship between the fluorescence intensity in each cell compartment was calculated using a linear regression model. All quantification data are from three biological replicates, and boxplots were generated using the ggplot2 function in RStudio (version 1.2.5033-1). Scripts for the analysis of imaging data have been publicly deposited in GitHub (https://github.com/AliceEseola/organelle-fluorescence-analysis-glm).

## Acknowledgments

This work was funded by a Halpin Scholarship in Rice Blast Research to A.B.E. and grants to NJT from the Gatsby Charitable Foundation and the Biotechnology and Biological Sciences Research Council (BBS/E/J/000PR9797, BB/V016342/1). M.J.E is supported by a National Science Foundation CAREER award (IOS-2141858).

## Notes

https://github.com/AliceEseola/organelle-fluorescence-analysis-glm)

### Competing Interest Statement

The authors have declared no competing interest.

