## Supplemental Table S2 for "Synchronous spatio-temporal control of autophagy and organelle trafficking is necessary for appressorium-mediated plant infection by *Magnaporthe oryzae*"

Supplementary Table S2: List of oligonucleotide primers used in this study

| **Primer Name** | **DNA Sequence (5'-3')** |
| --- | --- |
| pAE001-F | TGCAGCCCAATGTGGAATTCAAAAGCACAAAACTGAGTCACGGG |
| pAE001-R | CGAGGTGTACTCCTTCTGCAAAGA |
| pAE001.GFP-F | AAGGAGTACACCTCGATGGTGAGCAAGGGCGAGGAGCTG |
| pAE002-F | TGCAGCCCAATGTGGAATTCCTAGTAATACACGGTCACAAAGC |
| pAE002-R | GATAAATTCCCTCTGCTGGTAAAC |
| pAE002.GFP-F | CAGAGGGAATTTATCATGGTGAGCAAGGGCGAGGAGCTG |
| pAE003-F | TGCAGCCCAATGTGGAATTCCTTGATATCTGTGAATGACTCC |
| pAE003-R | AAGAGCCAGCTTGGTAGCA |
| pAE003.GFP-F | ACCAAGCTGGCTCTTATGGTGAGCAAGGGCGAGGAGCTG |
| pAE004-F | TGCAGCCCAATGTGGAATTCAGAGCGGAGTTGTTTATGGTGGTG |
| pAE004-R | ACCAGCATCCTCCCCTTCAGCCGC |
| pAE004.GFP-F | TGCAGCCCAATGTGGAATTCAATACTGGACATTAAGGGGAC |
| pAE005-F | TGCAGCCCAATGTGGAATTCCCTTATCGGTCTCAATCTCAACA |
| pAE005-R | TTGCCCACCCCTCCCCATAAACCT |
| pAE005.GFP-F | GGGAGGGGTGGGCAAATGGTGAGCAAGGGCGAGGAGCTG |
| pAE006-F | TGCAGCCCAATGTGGAATTCGTCATCACATTCAAGAACATT |
| pAE006-R | GGTGTTGCCATTCCTTTTCTT |
| pAE006.GFP-F | AGGAATGGCAACACCATGGTGAGCAAGGGCGAGGAGCTG |
| pAE007-F | TGCAGCCCAATGTGGAATTCTAGCAAGGAACAAAAGACAGGATC |
| pAE007-R | ATCGTACAAGCCCTCATCATCGCC |
| pAE007.GFP-F | GAGGGCTTGTACGATATGGTGAGCAAGGGCGAGGAG |
| pAE008.GFP-F | TCCTCCTTGGTTGTCTTGCATTGACCTTGGTGTTGCTGGCACAGGCG GTGAGCAAGGGCGAGGAGCTGTTC |
| pAE008.GFP-R | TTAGAGCTCGTCGTGCTTGTACAGCTCGTCCATGCCGTG |
| pAE008.p-F | TGCAGCCCAATGTGGAATTCTGTGATATCAACTTGGGGACT |
| pAE008.p-R | ACCAAGGTCAATGCAAGACAACCAAGGAGGATGACTAAAGACGGCAT CTTGACGGCGTTTTACTTTG |
| pAE008.trpC.T-F | CACGACGAGCTCTAAAGCGGCCGCCCGGCTGCAGCCCGG |
| GFP-F | ATGGTGAGCAAGGGCGAGGA |
| trpC-*Hin*dIII-R. | TCGACGGTATCGATAAGCTTAGTGGAGATGTGGAGTGGGCGCTT |

Underlines nucleotides are overhangs to the corresponding fragments
